# Nested oscillations and brain connectivity during sequential stages of feature-based attention

**DOI:** 10.1101/2020.02.28.969253

**Authors:** Mattia F. Pagnotta, David Pascucci, Gijs Plomp

## Abstract

Brain mechanisms of visual selective attention involve both local and network-level activity changes at specific oscillatory rhythms, but their interplay remains poorly explored. Here, we investigate anticipatory and reactive effects of feature-based attention using separate fMRI and EEG recordings, while participants attended to one of two spatially overlapping visual features (motion and orientation). We focused on EEG source analysis of local nested oscillations and on graph analysis of connectivity changes in a network of fMRI-defined regions of interest, and characterized a cascade of attentional effects and their interplay at multiple spatial scales. We discuss how the results may reconcile several theories of selective attention, by showing how β rhythms support anticipatory information routing through increased network efficiency and β-γ coupling in functionally specialized regions (V1 for orientation, V5 for motion), while reactive α-band desynchronization patterns and increased α-γ coupling in V1 and V5 mediate stimulus-evoked processing of task-relevant signals.

## 1. Introduction

Visual selective attention enables us to prioritize the processing of behaviorally relevant stimuli while filtering out irrelevant ones. This fundamental function of the human brain is supported by changes in activity patterns that occur at multiple spatial scales, from local neuronal circuits to global interactions between brain regions (Corbetta and Shulman, 2002; Greenberg et al., 2010; Kastner and Ungerleider, 2000; Scolari et al., 2014; Serences and Yantis, 2007). A central role in this distributed brain function is played by specific rhythms that coordinate selective processing both at the level of neuronal ensembles and at the macro-scale of multiple brain regions, determining the enhancement and propagation of attention-related signals or the downregulation of irrelevant activity (Antzoulatos and Miller, 2014; Bonnefond and Jensen, 2015; Buschman et al., 2012; Buzsáki and Draguhn, 2004; Canolty et al., 2007; Haegens et al., 2011a; Kopell et al., 2000; von Stein and Sarnthein, 2000).

A growing body of evidence suggests that distinct neuronal rhythms may serve distinct functional roles in selective attention. Activity over an extended frequency range around the alpha rhythm (α, 7– 14 Hz), for instance, has been widely linked to the prevention or inhibition of task-irrelevant signals (Chelazzi et al., 2019; Foster and Awh, 2019; Van Diepen et al., 2019; but see Noonan et al., 2016; Schroeder et al., 2018). A potential mechanism by which α rhythms would gate selective attention is through the pulsed inhibition of ongoing cortical activity, by providing phasic bursts of inhibition that suppress the functional processing and inter-areal communication in the gamma frequency range (γ, above 30 Hz) (Jensen and Mazaheri, 2010). In support of this, several studies have recently documented an inverse relationship between α-band feedback signaling from attentional control regions and feedforward γ-band activity in sensory cortices, revealing top-down, attention-related α-modulations that interfere with local structures of α-γ phase-amplitude coupling (PAC) in sensory areas (Bonnefond and Jensen, 2015; Haegens et al., 2011b; Mathewson et al., 2011; Mazaheri and Jensen, 2010; Pascucci et al., 2018; Popov et al., 2017). At a relatively higher range of frequencies, beta rhythms (β, 15–30 Hz) have been instead related to the instantiation of distributed task-relevant representations that convey context- and content-specific information (Antzoulatos and Miller, 2016, 2014; Buschman et al., 2012; Richter et al., 2018; Spitzer and Haegens, 2017). At the same time, β-band activity has been also shown to promote feedforward and inter-areal γ-band synchronization (Richter et al., 2017), while preventing interference and irrelevant attentional shifts (Fiebelkorn and Kastner, 2019). Thus, distinct rhythms appear to fulfil separate but complementary roles in selective attention, by differentially modulating local activity, structures of cross-frequency coupling and neuronal communication in large-scale networks.

A further distinction exists as to whether specialized rhythms support the anticipatory (pre-stimulus) and reactive (post-stimulus) components of selective attention. Pre-stimulus synchronization of α-band activity has been mostly described in relation to anticipatory, proactive-like mechanisms to filter-out irrelevant features and locations (Foxe and Snyder, 2011; Kelly et al., 2006; Oliveira et al., 2014; Snyder and Foxe, 2010; Worden et al., 2000; but see Noonan et al., 2016), whereas post-stimulus, α-band event-related desynchronization (ERD) has been linked to the release from inhibition (Klimesch et al., 2007), which in turn facilitates local and feedforward γ-band activity conveying task-relevant signals (Bonnefond et al., 2017; Pascucci et al., 2018; Popov et al., 2017). Conversely, post-stimulus increases of α-band activity may represent reactive suppression and the return to inhibitory states that prevent task-irrelevant processing (Pascucci et al., 2018). In a similar vein, increases of β-band activity have also been found in preparatory stages of processing (Schneider and Rose, 2016), as top-down modulatory signals that relay task-specific behavioral context (Richter et al., 2018), or in more reactive stages as feedback control mechanisms that facilitate the bottom-up communication of attended stimuli (Bastos et al., 2015). Crucially, most of these phenomena have been investigated separately and the exact nature of their interplay, as well as the complexity of local and large-scale neuronal dynamics involved in selective attention, remains poorly understood.

In the present work, we leveraged different spatial scales of neuronal activity to characterize the temporal dynamics of local and network-level changes during anticipatory and reactive stages of attentional processing. At the local level, we focused on the phenomenon of nested oscillations (Bonnefond et al., 2017), hypothesizing functional differences in the coupling between α-β and γ rhythms depending on the anticipation and processing of relevant or irrelevant sensory signals. At the network level, we exploited measures of directed connectivity among distributed attentional and sensory areas, with the goal to relate large-scale functional interactions to local changes in nested oscillations.

To this end, we acquired separate EEG and fMRI recordings while human participants were performing basic visual tasks under different feature-based attention conditions. We presented two spatially overlapping features (Baldauf and Desimone, 2014), motion and orientation, and we manipulated participants attention toward each of them in separate blocks, probing both anticipatory and reactive feature-specific attentional responses. Our results revealed a cascade of attentional effects at multiple spatial scales, from anticipatory β-band coupling and connectivity changes involving functionally specialized regions (e.g., V1 and V5), to distinct local and global post-stimulus dynamics initiated by α-band desynchronization.

## 2. Results

### 2.1. Attention modulates visual-evoked potentials

The behavioral task required the participants to attend visual stimuli presented at central location (Figure 1A), and to discriminate either the motion direction of signal-dots in the Random Dot Kinematograms (RDK; motion discrimination task), or the off-vertical tilt of the Gabor (orientation discrimination task). Hereafter, these two tasks conditions are named Attended-motion and Attended-orientation, respectively. In both Attended conditions, the stimuli contained both features (motion and orientation). In a third task condition that served as control, participants were asked to report sporadic color changes in the fixation spot, which rendered the same stimuli irrelevant for the task (Unattended condition), without changing their physical characteristics. We calibrated task-difficulty beforehand (see Methods) and did not observe differences in behavioral performance between the two discrimination tasks. Percentage correct was 89.0% (*SD*=5.2) and 88.6% (*SD*=7.2) for Attended-motion and Attended-orientation respectively (*t*=0.19, *p*=0.8485), with corresponding reaction times (RTs) of 687.5 ms (*SD*=159.1) and *M*=699.5 ms (*SD*=141.0; *t*=-0.62, *p*=0.5432). In the Unattended control condition, participants correctly detected a color change in the fixation spot in 99.2% of target trials (*SD*=1.4), with RT of 484.4 ms (*SD*=46.6).

**Figure 1.**
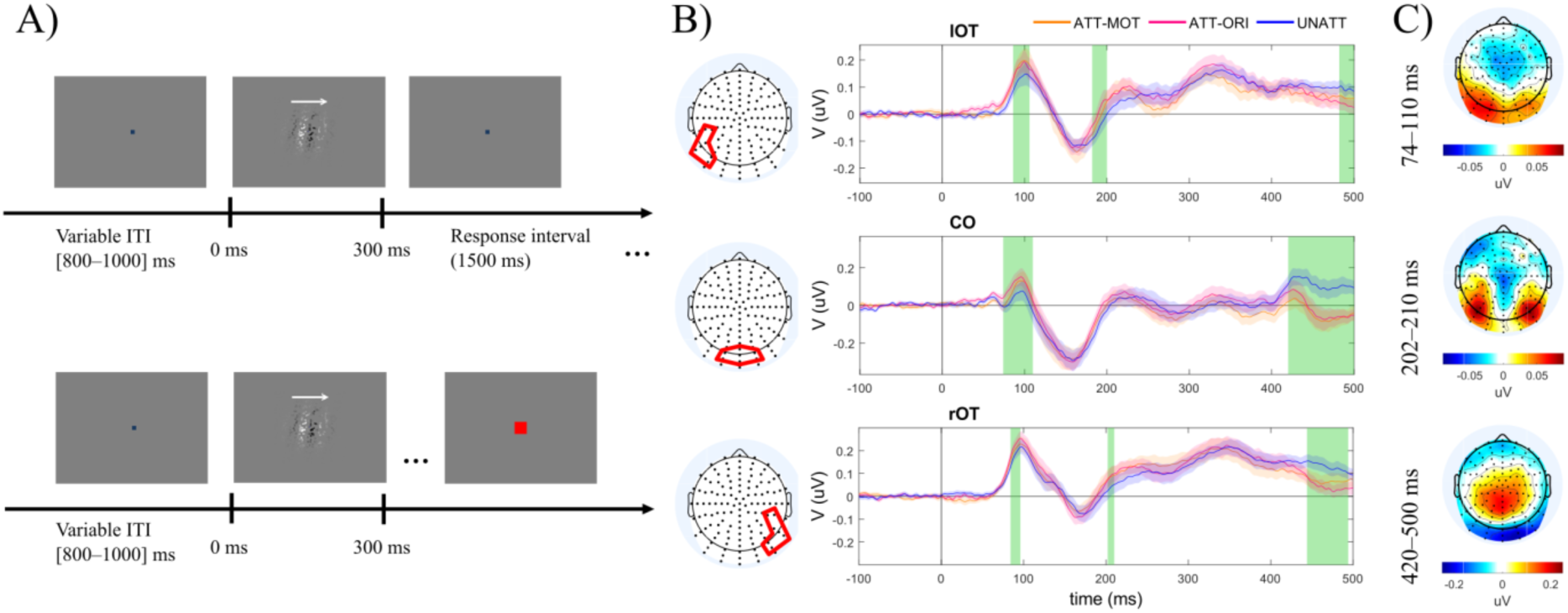
Experimental paradigm and VEPs results. A) Schematic representation of trial structures for Attended (motion or orientation discrimination) and Unattended (detection of color changes in the fixation spot). The white arrows on top of the visual stimuli indicate the direction of coherent motion and were not present during the experiment. The red fixation spot is enlarged for illustrative purpose, it did not change size during color changes. B) Show grand-average VEPs from three electrode clusters: left occipito-temporal (lOT); centro-occipital (CO); right occipito-temporal (rOT). Electrode locations of each cluster are highlighted in red in the corresponding scalp representation (on the left). The VEPs in the Attended-motion (ATT-MOT) and Attended-orientation (ATT-ORI) conditions are shown in orange and purple, respectively; while, VEPs in the Unattended condition (UNATT) are shown in blue. For each condition, the shading represents the standard error of the mean. Green vertical shades represent time intervals that showed statistically significant results in the repeated-measures ANOVA (*p*_*FDR*_<0.05). Results of repeated-measures ANOVA and of post-hoc analyses are provided in Table 1. C) The figures show the scalp topographical distribution of the differences between Attended-motion and Unattended, at the time intervals derived from the ANOVA results (top to bottom).

Attention has been shown to modulate the amplitude of stimulus-evoked P1 component, at around 100 ms post-stimulus onset (Hillyard et al., 1998; Pascucci et al., 2018; Zhang and Luck, 2009). Comparing Attended conditions with the Unattended showed an attentional P1 increase over posterior electrodes (Figure 1B). Post-hoc analyses revealed significant P1 amplitude increases for both Attended-motion and Attended-orientation in left occipito-temporal (lOT) and centro-occipital (CO) electrode clusters. In right occipito-temporal (rOT) electrodes, P1 amplitude was significantly higher than Unattended only for Attended-orientation (Table 1).

**Table 1.**
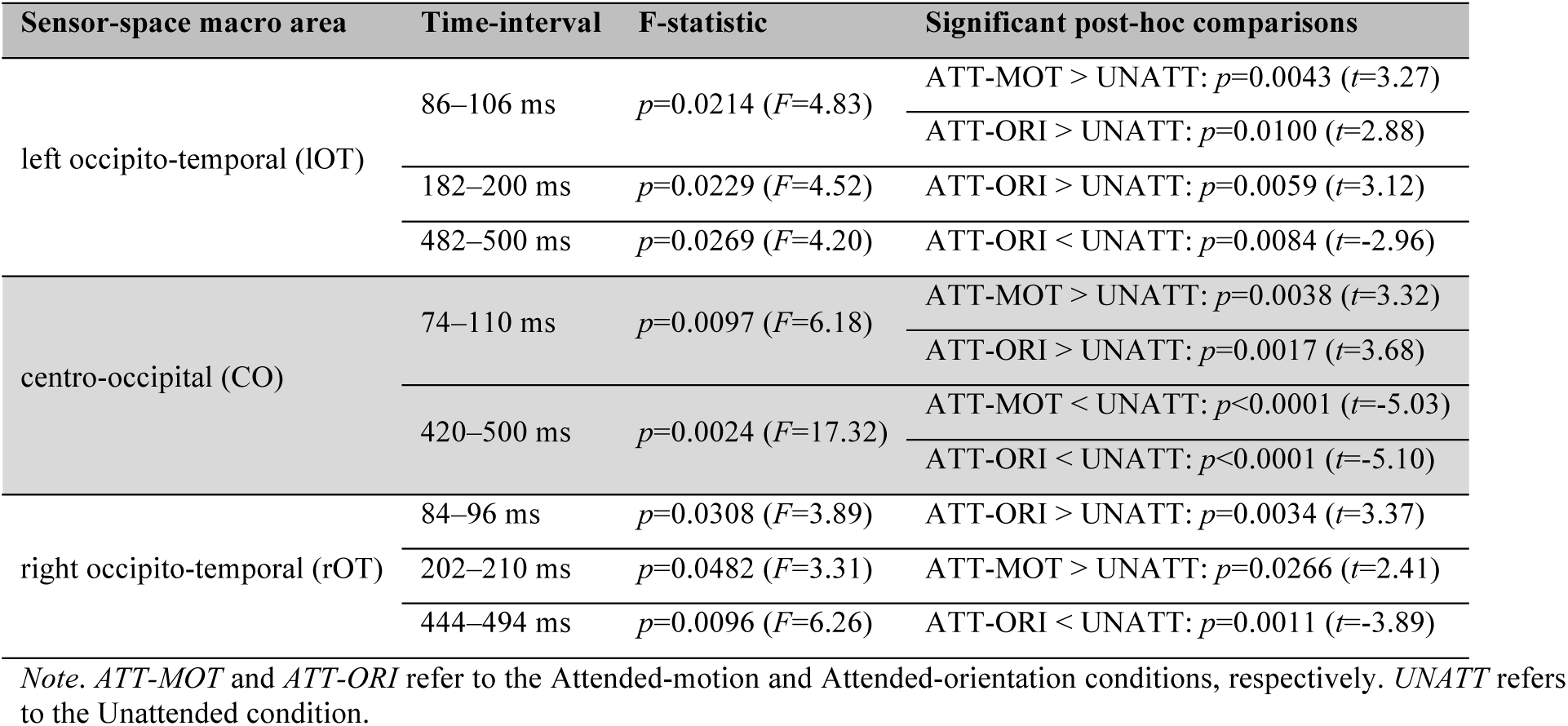
VEPs analysis; results of repeated measures ANOVA and post-hoc tests.

The results further showed attentional modulations of the P2 component (around 200 ms after stimulus onset), with significantly increased amplitudes for Attended-orientation in lOT (*p*=0.0059), and for Attended-motion in rOT (*p*=0.0266). The effects replicate previous findings of P2 modulations for target stimuli (Kanske et al., 2011; Maeno et al., 2004; Van Voorhis and Hillyard, 1977). At latencies beyond 400 ms, significant differences in VEPs were mostly observed over CO, with decreased amplitude for both Attended conditions (Figure 1B, Table 1). The sign and topographical distribution of these late differences resembled the posterior selection negativity (Figure 1C) (Hillyard and Anllo-Vento, 1998). While selection negativity is usually observed from 150-200 ms after stimulus onset with static stimuli (Daffner et al., 2012; Hillyard and Anllo-Vento, 1998; Pascucci et al., 2018), our use of moving stimuli with multiple features may explain the late occurrence.

### 2.2. Attention dynamically modulates brain rhythms

To investigate how attention modulates brain rhythms before and after stimulus onset, we compared spectral power distributions between attention conditions. Across the pre-stimulus interval (−300–0 ms), we found that attention significantly enhanced power spectra. The anticipatory increases for the Attended conditions were most prominent in the β-band and distributed over lateral and occipito-temporal electrodes (Figure 2A). To localize these anticipatory effects to underlying cortical areas, we next performed the power-spectral analysis on source-reconstructed signals. We used EEG source imaging based on individual lead-fields to extract the signals from 22 cortical regions of interest (ROIs) across the brain (Oostenveld et al., 2011; Rubega et al., 2019; Van Veen et al., 1997). The ROIs were derived from fMRI cluster-peaks of significant differences, obtained with the same tasks and participants (Figure 3, Table 2) (see Methods). The EEG source-space results showed that the anticipatory β-band power increases in Attended conditions mainly originate from lateral visual areas (Figure 2B).

**Table 2.** Details about the 22 regions of interest (ROIs).

| Anatomical region | ROI abbreviation | Coordinates of centroid (MNI-space) |  |  |
| --- | --- | --- | --- | --- |
| | | $x$ | $y$ | $z$ |
| left premotor area | PM L | -43.60 | -6.40 | 41.20 |
| right premotor area | PM R | 42.61 | -6.00 | 47.13 |
| left middle frontal gyrus | MFG L | -32.00 | 30.00 | 42.00 |
| right middle frontal gyrus | MFG R | 34.29 | 28.29 | 42.29 |
| left inferior frontal gyrus | IFG L | -45.00 | 27.13 | 6.75 |
| right inferior frontal gyrus | IFG R | 44.29 | 18.29 | 25.14 |
| supplementary motor area | SMA | -2.50 | -0.17 | 59.67 |
| middle cingulate cortex | MCC | 10.00 | -24.00 | 42.00 |
| left calcarine sulcus | V1 L | -4.29 | -93.14 | 2.29 |
| right calcarine sulcus | V1 R | 12.00 | -92.00 | 2.00 |
| left cuneus | Cuneus L | -6.67 | -81.00 | 26.33 |
| left secondary visual cortex | V2 L | -36.47 | -83.06 | -3.76 |
| right secondary visual cortex | V2 R | 35.00 | -86.00 | -7.00 |
| left fusiform gyrus | FFG L | -35.00 | -62.67 | -13.67 |
| left gyrus postcentralis | GPC L | -33.33 | -36.00 | 51.33 |
| left superior parietal lobule | SPL L | -10.00 | -50.29 | 45.14 |
| right superior parietal lobule | SPL R | 18.00 | -63.50 | 52.50 |
| left inferior parietal lobule | IPL L | -55.33 | -34.67 | 32.67 |
| right inferior parietal lobule | IPL R | 49.41 | -48.94 | 37.41 |
| left middle temporal area | MTG L | -55.00 | -56.00 | 13.00 |
| right visual cortex V5 | V5 R | 53.33 | -61.00 | 1.33 |
| right middle temporal area | MTG R | 57.50 | -37.25 | -5.75 |

*Note.* In the column about 'ROI abbreviation': $L$ and $R$ stand for left and right, respectively.

**Figure 2.**
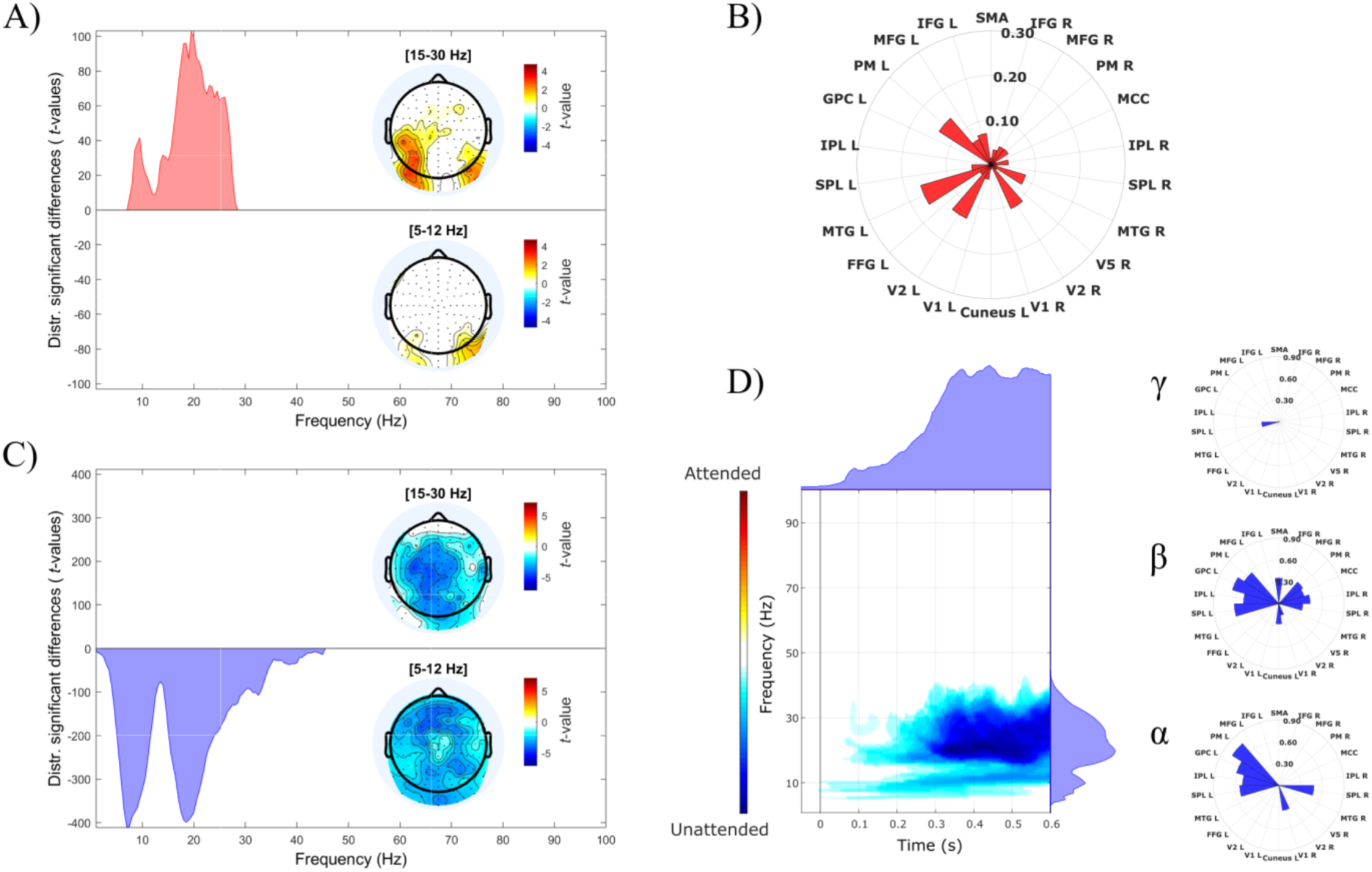
Attentional modulation of brain rhythms: comparison between Attended-motion and Unattended. Results of sensor-space power spectra analysis are shown for anticipatory (A) and reactive (C) time intervals. Each figure represents the distribution of statistically significant differences (positive *t*-values for Attended-motion higher than Unattended, and vice versa negative *t*-values) from cluster-based permutation (two-tailed *t*-test with *p*<0.05, 50000 permutations, and *p*<0.05 for the permutation test), together with the topographical distributions of these differences in the α-band (bottom) and β-band (top). Here and in the following, red indicate positive differences between Attended and Unattended conditions (i.e., increases with attention), while blue indicates negative differences (i.e., decreases with attention). B) The polar histogram represents the effect sizes for each ROI of source-space power differences between conditions in the β-band (15–30 Hz). D) The figure shows the results of source-space time-varying power spectra analysis, with the distribution of statistically significant differences from cluster-based permutation (two-tailed *t*-test with *p*<0.05, 50000 permutations, and *p*<0.05 for the permutation test). The marginal plots show time- or frequency-collapsed distributions of significant differences in power spectra. The polar histograms on the right represent the effect sizes for each ROI of power differences between conditions, in the following frequency bands: α (5–12 Hz), β (15–30 Hz), and γ (45–100 Hz). Effect sizes were estimated using Cohen’s *d* (Cohen, 1992). Analogous results were obtained when comparing Attended-orientation to Unattended (Figure S1).

**Figure 3.**
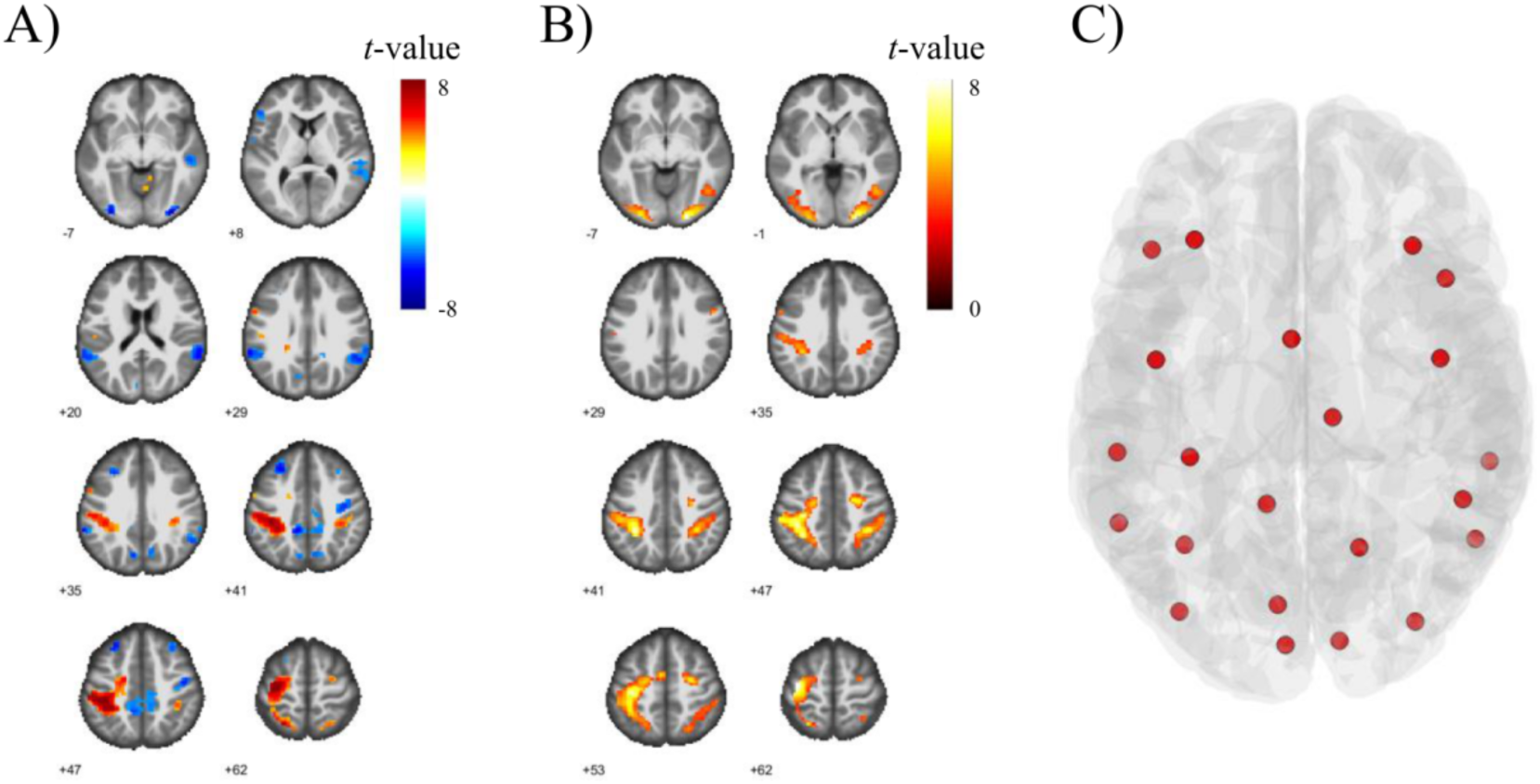
fMRI results and ROIs. A) Results of the voxel-wise *t*-test comparison across participants for the contrast Attended vs. Unattended; positive and negative results from the comparison are shown in the red and blue color scales, respectively. B) Results of the voxel-wise *t*-test comparison across participants for the contrast Attended vs. Rest; voxels with higher activity in both Attended conditions compared to Rest are shown in the yellow color scale. C) Centroids of the 22 ROIs identified from the fMRI results.

In the post-stimulus interval, we found greater decreases in α and β-band power for the Attended conditions across widespread electrodes (Figure 2C). These effects reflect an ERD (Minami et al., 2014; Pfurtscheller and Lopes da Silva, 1999; Schmiedt et al., 2014; Yordanova et al., 2001) that was larger for attended than unattended stimuli (Mazaheri and Picton, 2005; Pascucci et al., 2018). We used time-varying spectral analysis in the source-space and showed that significant differences between Attended and Unattended conditions emerged shortly after stimulus onset, with stronger ERD in the α and β-band for Attended in fronto-parietal regions (Figure 2D).

Taken together, the results showed that attention modulates local brain rhythms in distinct ways before and after stimulus onset. Anticipatory effects involved attentional increase of β-band power in lateral visual areas, while reactive effects involved stronger α and β-band ERD in parietal and frontal regions due to attention. Spectral power changes occurred irrespective of the attended feature (motion or orientation). These results suggest a dynamic reorganization of distributed activity in distinct frequency bands, depending on whether a stimulus is attended or not. We further investigated the pre- and post-stimulus effects of attentional processing using network and PAC analyses.

### 2.3. Anticipatory stage: increased large-scale network efficiency

To identify anticipatory network changes due to attention, we estimated the frequency-specific directed connections between the 22 ROIs using a multivariate functional connectivity measure, the information partial directed coherence (iPDC) (Baccalá and Sameshima, 2014; Takahashi et al., 2010). To describe and compare the topological properties of the resulting networks in different frequencies, we used two network measures: global efficiency and local efficiency (Latora and Marchiori, 2001; Rubinov and Sporns, 2010). Given the ability of the iPDC to characterize both directionality and directness of inter-areal connections and because of its information-theoretic foundation (Baccalá and Sameshima, 2014; Takahashi et al., 2010), these functional connections can be seen as paths of integration between network’s ROIs. The iPDC-derived network’s global efficiency represents the frequency-specific level of global functional integration among all regions. Likewise, the network’s local efficiency represents the level of functional integration within subgraphs of neighbor regions, defined by strongest functional connections, while the local efficiency of each single ROI measures the local integration within its own subgraph only (see Methods).

Topological network analysis revealed that attention to motion significantly increased both global and local efficiency of the network in the β (15–21 Hz) and high-γ band (around 60 Hz) (Figure 4A). We found analogous increases for Attended-orientation (Figure 4B), where statistically significant differences in the γ-band spanned a larger frequency range (57–71 Hz) and we observed an additional increase in global efficiency at 33-34 Hz. The results suggest that anticipatory network effects could enhance large-scale network communication and information routing in the β and γ-band, by changing network topology at these frequencies to favor functional integration both globally and within functional subgraphs.

**Figure 4.**
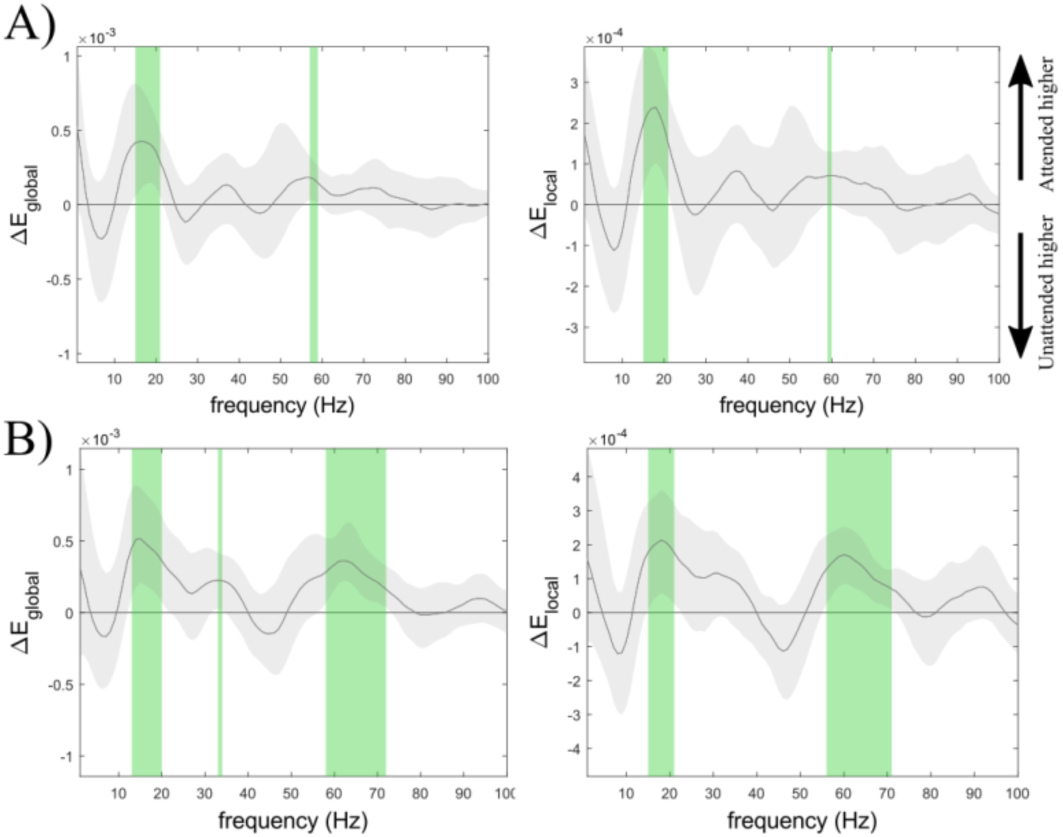
Anticipatory differences in network efficiency. Results for network’s global efficiency and local efficiency, for the comparisons: A) Attended-motion vs. Unattended; B) Attended-orientation vs. Unattended. Differences between conditions are evaluated as a function of frequency for global (left) and local efficiency (right). Gray shading represents the standard error of the mean. Green vertical shades represent the frequencies that showed statistically significant results in a two-tailed *t*-test (*p*_*FDR*_<0.05).

### 2.4. Anticipatory stage: local changes in β-γ coupling

Before stimulus onset we found that attention enhanced network functional integration in the β and γ-band. The activity from the two frequency bands may be integrated locally through mechanisms of cross-frequency coupling (Bonnefond et al., 2017; Buzsáki, 2006; Canolty et al., 2006; Canolty and Knight, 2010; Jensen and Colgin, 2007), we thus investigated whether attention modulates the within-region PAC (Martínez-Cancino et al., 2019; Penny et al., 2008). For this analysis, we selected 9 ROIs in occipito-temporal cortex based on the results of power spectra analyses, and in line with previous studies that reported modulations of low-frequency activity and cross-frequency coupling in occipital cortex (Bastos et al., 2015; Bonnefond and Jensen, 2015; Pascucci et al., 2018; Schmiedt et al., 2014). The results showed that attention to motion selectively increased β-γ PAC in region V5-R (*f*_*PHASE*_: 24– 27 Hz; *f*_*AMPLITUDE*_: 54–80 Hz) (Figure 5A). For Attended-orientation, we found a significant β-γ PAC increase at high amplitude frequencies (*f*_*PHASE*_: 23–25 Hz; *f*_*AMPLITUDE*_: 76–86 Hz) and decrease at lower frequencies (*f*_*PHASE*_: 19–20 Hz; *f*_*AMPLITUDE*_: 44–56 Hz) in region V1-L, and increased PAC in region MTG-L (*f*_*PHASE*_: 20–22 Hz; *f*_*AMPLITUDE*_: 46–62 Hz) (Figure 5B). Furthermore, we observed a significant β-γ PAC increase in region V2-R and decrease in region V2-L, for both Attended conditions compared to Unattended. The anticipatory PAC effects could not be attributed to power differences in the signals, as shown by a control analysis based on stratification (Aru et al., 2015; Oostenveld et al., 2011) (Figure S2). In sum, while certain regions showed general attentional modulations of PAC (V2-L, V2-R), others showed effects that were specific of the attended feature: V5-R for motion, V1-L and MTG-L for orientation.

**Figure 5.**
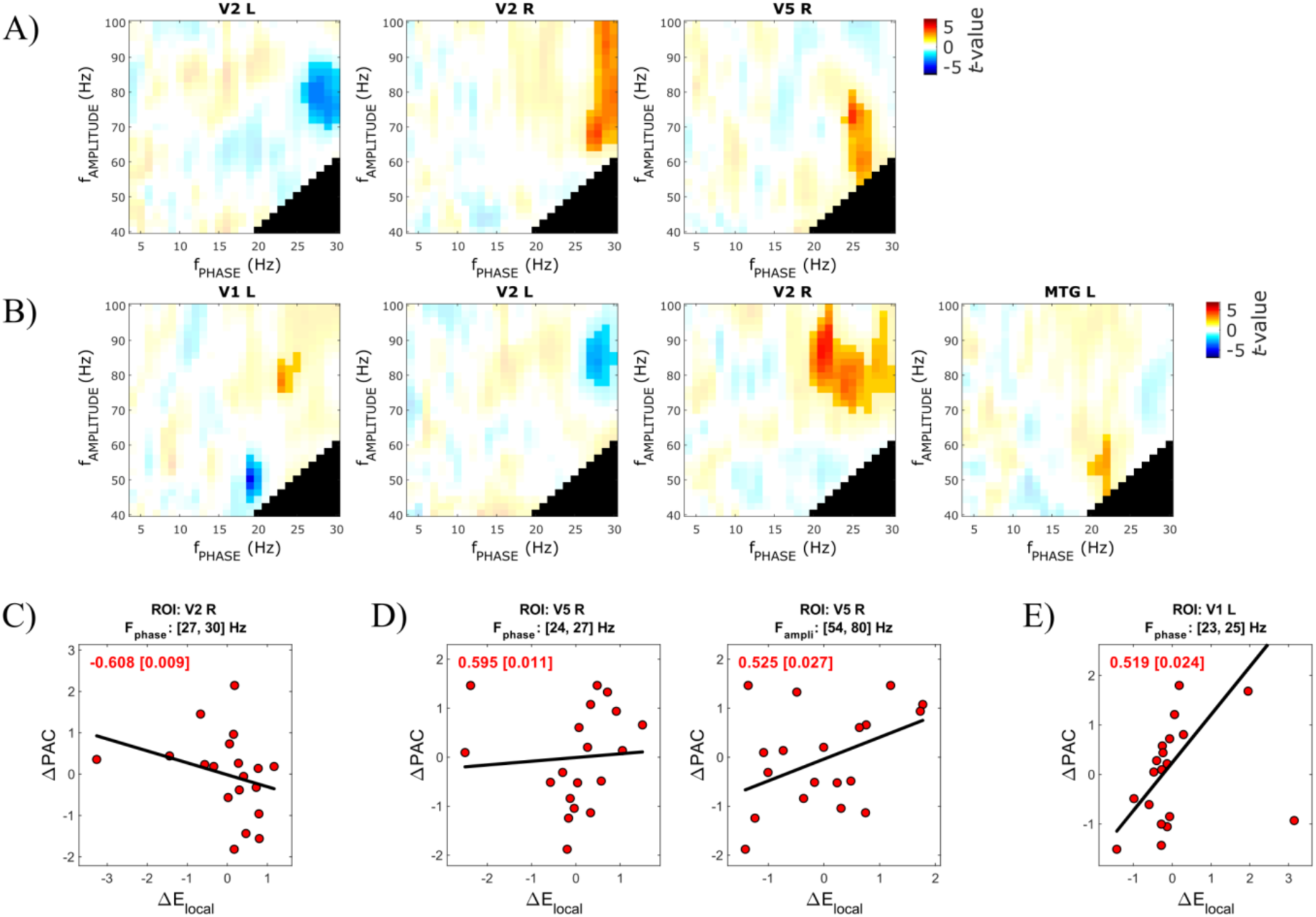
Anticipatory differences in local PAC and correlation with network’s local changes. Differences in within-region anticipatory PAC are shown for each pair *f*_*PHASE*_– *f*_*AMPLITUDE*_, for Attended-motion compared to Unattended (A) and for Attended-orientation compared to Unattended (B). The figure shows regions with significant differences from the cluster-based permutation comparison (two-tailed *t*-test with *p*<0.05, 50000 permutations, and *p*<0.05 for the permutation test), which are highlighted as more opaque. Black areas indicate *f*_*PHASE*_–*f*_*AMPLITUDE*_ pairs excluded from the analysis (see Methods). Figures C–E report the within-region correlation between variations in PAC (*ΔPAC*) and variations in single-node’s local efficiency (*ΔE*_*local*_), for regions that showed a significant correlation (*p*<0.05): C) V2-R for Attended-motion; D) V5-R for Attended-motion; E) V1-L for Attended-orientation. Each figure reports the Spearman’s *rho* and the *p*-value for testing the null-hypothesis of no correlation (inside brackets).

### 2.5. A link between PAC and local efficiency modulations

Before stimulus onset, we found both enhanced large-scale network integration in the β and γ-band (Figure 4) and increased β-γ PAC in specific occipito-temporal regions (Figure 5A–B). Therefore, we next asked whether in these regions there is a linear relation between the local cross-frequency coupling and their level of functional integration within the network. In each ROI, we evaluated the relationship between changes of PAC (*ΔPAC*) and changes of its local efficiency (*ΔE*_*local*_) due to attention (see Methods). The results showed a significant negative correlation in region V2-R between β-γ *ΔPAC* and β-band *ΔE*_*local*_ for the comparison between Attended-motion and Unattended (*rho*=-0.608, *p*=0.009, number of outliers = 1) (Figure 5C), which means that when β-band local efficiency increased with attention, the local β-γ PAC decreased. For the same comparison, we found a reversed relationship in region V5-R, where β-γ *ΔPAC* correlated positively to both β-band *ΔE*_*local*_ (*rho*=0.595, *p*=0.011, number of outliers = 1) and γ-band *ΔE*_*local*_ (*rho*=0.525, *p*=0.027, number of outliers = 1) (Figure 5D). These effects reveal that in V5-R when local efficiency increased with attention in the β and γ-band, the local β-γ PAC increased too, across subjects. From comparing Attended-orientation and Unattended, a significant positive correlation between β-γ *ΔPAC* and β-band *ΔE*_*local*_ was observed in region V1-L (*rho*=0.519, *p*=0.024, number of outliers = 0) (Figure 5E). No statistically significant effects were observed in the other ROIs tested.

It is worth highlighting that a significant positive correlation between β-γ *ΔPAC* and β-band *ΔE*_*local*_ was found in V5-R when motion was attended and in early visual area V1-L when orientation was attended, which are respectively known to be sensitive to motion coherence and Gabor’s orientation (Ahlfors et al., 1999; Bosking et al., 1997; Koelewijn et al., 2011; Schoenfeld et al., 2007; Simoncelli and Olshausen, 2001). The effects therefore suggest that, only in functionally specialized regions, the anticipatory within-region increases of local cross-frequency coupling are linked to the increased levels of functional integration among neighboring areas of that region.

### 2.6. Reactive stage: dynamic local changes in α-γ coupling

We found larger stimulus-evoked α and β-band desynchronization for attended stimuli (Figure 2). We therefore investigated whether α-γ PAC and β-γ PAC were modulated by reactive attentional processing. Our results showed significantly increased α-γ PAC for Attended-motion in regions V5-R and MTG-R (Figure 6A). The α-γ PAC increase in V5-R started around 46 ms post-stimulus onset and lasted up to approximately 100 ms, involving high γ-band frequencies (64–100 Hz). The significant α-γ PAC increase in MTG-R occurred between 66 and 124 ms, modulating frequencies in the range 72– 100 Hz. For Attended-orientation, we found that α-γ PAC in V1-L was significantly increased compared to Unattended, at latencies and frequencies in the ranges 86–130 ms and 62–100 Hz, respectively (Figure 6B). A recent study showed a similar reactive increase in α-γ PAC using an orientation discrimination task (Pascucci et al., 2018). No statistically significant effects were observed in the other ROIs tested, and no correlations between changes of within-region α-γ PAC and local efficiency were observed in post-stimulus time.

**Figure 6.**
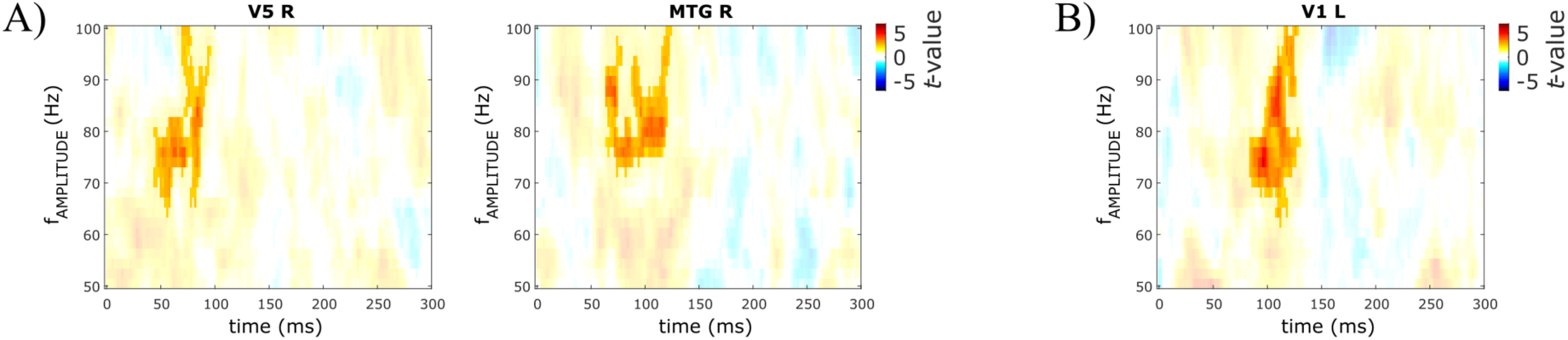
Stimulus-evoked differences in local α-γ PAC. The temporal evolution of differences in PAC are shown for the coupling between α-band phase (10 Hz for Attended-motion compared to Unattended, and 8 Hz for Attended-orientation compared to Unattended) and γ-band amplitude (50–100 Hz, in 2Hz-steps). Panel A) shows Attended-motion compared to Unattended, B) Attended-orientation compared to Unattended. Result were obtained via cluster-based permutation (two-tailed *t*-test with *p*<0.05, 50000 permutations, and *p*<0.05 for the permutation test), statistically significant results are highlighted as more opaque.

It should be noted that the reactive attentional increases in α-γ PAC involved the same functionally specialized regions that showed anticipatory increases in β-γ PAC (V5-R for Attended-motion and V1-L for Attended-orientation), positively correlated at between-subject level with β-band *ΔE*_*local*_ (Figure 5). We did not find, however, any significant differences in stimulus-evoked β-γ PAC between Attended and Unattended conditions. Taken together, this suggests that the anticipatory and reactive stages of attentional processing may be distinctively mediated by β and α rhythms.

### 2.7. Reactive stage: fast dynamics of network efficiency

We next asked how attentional processing affects large-scale network topology after stimulus onset. To dynamically investigate stimulus-evoked network changes, we employed time- and frequency-resolved estimates of functional connectivity (iPDC) between the 22 ROIs (Pascucci et al., 2019; Takahashi et al., 2010). The results showed significantly diminished global and local efficiency of the network for Attended-motion compared to Unattended in both α and β-band, and from latencies around 270 ms and 300 ms, respectively (Figure 7). These low-frequency differences in network topology were preceded by a significant increase for Attended-motion of both global and local efficiency in the γ-band, at around 160 ms post-stimulus onset (Figure 7C). Similar results were obtained by comparing Attended-orientation and Unattended, where we found significantly decreased global and local efficiency of the network at low-frequencies (α and β-band), roughly from 240 ms post-stimulus onset, preceded by a significant γ-band increase for Attended-orientation around 180 ms post-stimulus onset in both efficiency measures (Figure S3).

**Figure 7.**
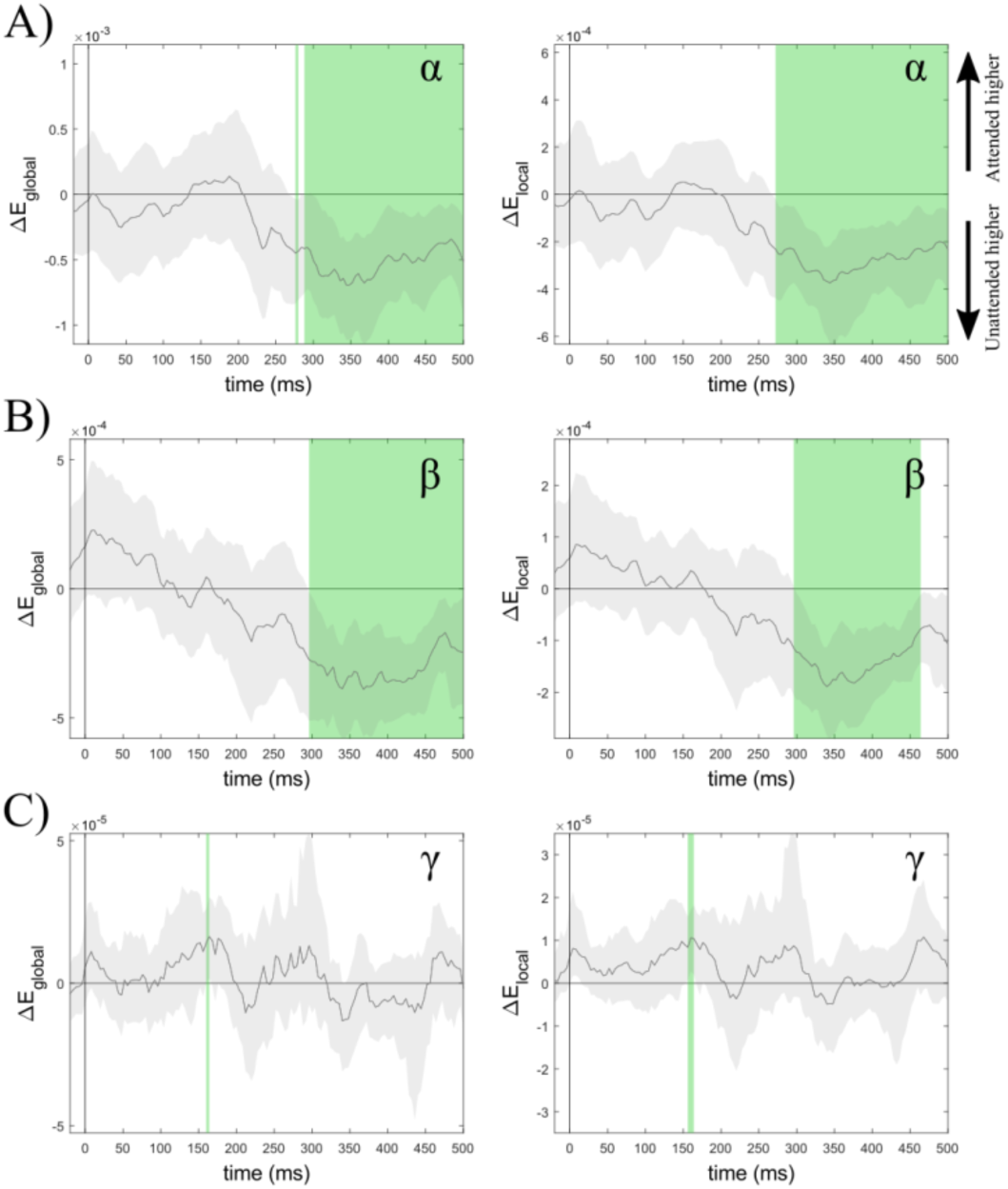
Stimulus-evoked differences in network efficiency. Differences between Attended-motion and Unattended are shown for three frequency bands, A) α-band, B) β-band, and C) γ-band. Left panels show network’s global efficiency, right panels show network’s local efficiency. Gray shading represents the standard error of the mean. Green vertical shades represent latencies that showed statistically significant results in the two-tailed *t*-test (*p*_*FDR*_<0.05).

Reactive network changes occurred in rapid succession: attention first enhanced efficiency measures in the γ-band (∼150–200 ms) and then reduced them in the α and β-band (after 240 ms post-stimulus onset). The low-frequency effects at longer latencies indicate that the evoked α and β-band desynchronization (Figure 2) co-occurs with network-level changes that reduce the functional integration both globally and within subgraphs, favoring more local processing.

## 3. Discussion

In the present work, we sought to characterize local and large-scale neuronal dynamics during anticipatory and evoked stages of feature-based selective attention. To this aim, we investigated local cross-frequency coupling and brain-wide directed connectivity during a visual discrimination task that required the attentional selection of one of two spatially overlapping features. Our results show distinct patterns of nested oscillations and functional network topologies that precede and follow stimulus onset. More precisely, we found evidence for a dominant role of β rhythms in the anticipatory stage, followed by stimulus-evoked α-band desynchronization patterns and α-driven structures of nested oscillations. The use of a control condition and the presentation of overlapping features at a single location also allowed us to disentangle between general effects of stimulus relevance and specific effects of the attended feature (motion or orientation). Figure 8 summarizes the findings, highlighting the sequential evolution of local and network-level phenomena with respect to stimulus onset.

**Figure 8.**
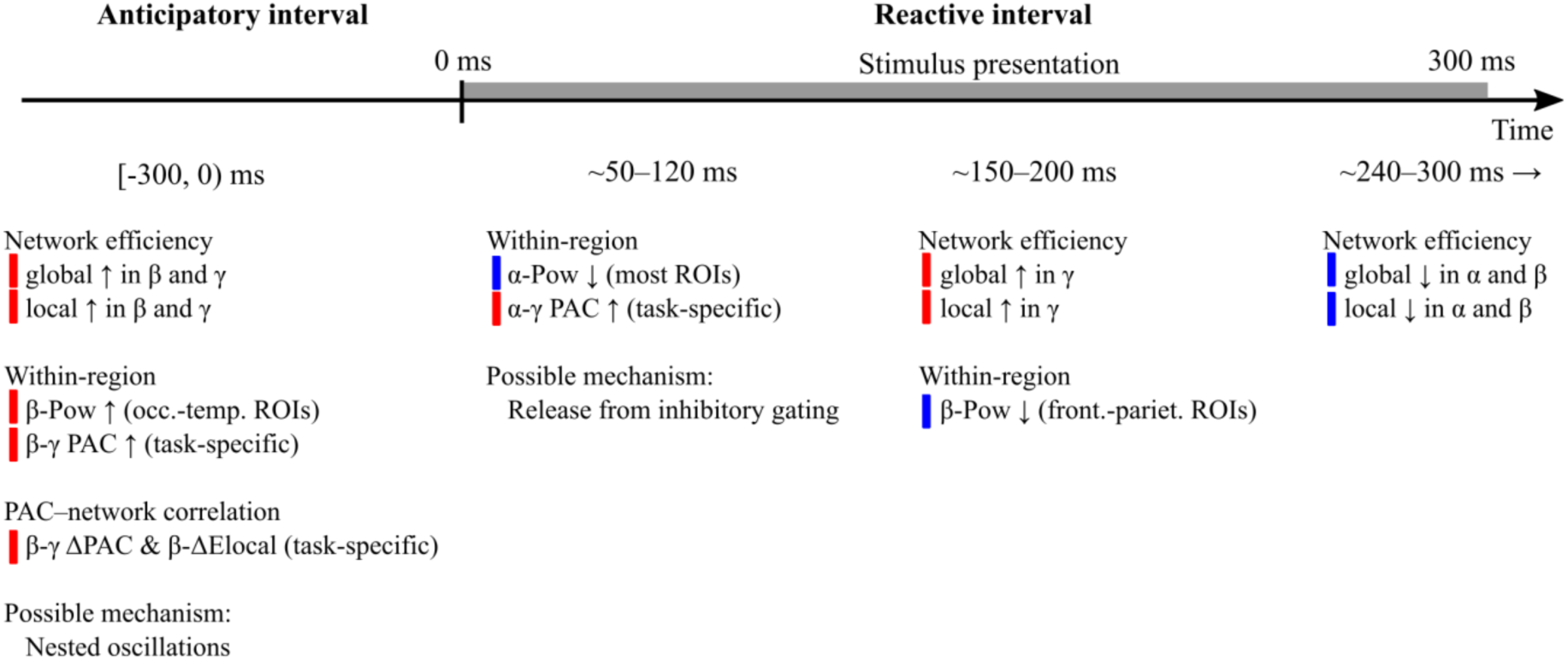
Temporal sequence of the effects induced by visual selective attention. Red (blue) patches indicate attentional increases (decreases), as do upwards (↑) and downwards (↓) arrows. The abbreviation *Pow* refers to the spectral power.

As evident from Figure 8, feature-based selective attention was mediated by a stimulus-induced reconfiguration of both local and large-scale neuronal interactions. Before stimulus onset, increased levels of network integration in the β and γ-band were paralleled by local increases of β-γ coupling in occipito-parietal regions, and the two phenomena correlated across participants. After stimulus onset, brain activity underwent a marked reconfiguration, with robust desynchronization of α rhythms and increased α-γ coupling in feature-selective regions (V5 and MTG for motion discrimination, V1 for orientation discrimination).

The observed cascade of attention-related effects suggests that different mechanisms, involving the interplay between large-scale dynamics and nested oscillations, serve distinct aspects of selective attention. Pre-stimulus β-band activity may reflect an increase in functional communication among regions involved in the upcoming task, with a consequent boost of high-frequency activity in functionally specialized areas (Buzsáki and Wang, 2012). Interpreting the role of β rhythms in the context of attention and cognitive processing, however, is not trivial, given the large body of work traditionally relating these rhythms to motor control (Pfurtscheller et al., 1996; Pogosyan et al., 2009; Sanes and Donoghue, 1993; Swann et al., 2009), sensorimotor functions (Lalo et al., 2007) and working memory (Lundqvist et al., 2016; Schneider and Rose, 2016; Siegel et al., 2009). A recent line of evidence has associated β rhythms with the maintenance of the status quo, that is, the preservation of ongoing sensorimotor states and cognitive sets (Engel and Fries, 2010). Following this line, other studies have hypothesized a role for β-band synchronization in establishing flexible networks of neuronal ensembles that carry content- and task-specific representations (Antzoulatos and Miller, 2016, 2014; Buschman et al., 2012; Spitzer and Haegens, 2017). Our results appear in agreement with this latter definition and extend it, by showing that large-scale network interactions associated with β rhythms may control endogenous mechanisms of attention before stimulus onset, likely conveying task-related information to downstream areas (e.g., the type of stimulus to attend/ignore, the type of response required) (Richter et al., 2017).

It is relevant to note that, contrary to previous reports (see Chelazzi et al., 2019; Foster and Awh, 2019; Snyder and Foxe, 2010 for reviews and discussions), we found no evidence of anticipatory modulations of α-band activity in our data. While, at first, this seems to support the alternative view of α rhythms as a secondary type of inhibitory mechanism, observable only in relation to target processing (e.g., at ipsilateral sides of relevant stimuli; Chelazzi et al., 2019; Foster and Awh, 2019), the fact that our analysis always involved subtracting a control condition with identical stimulation could have hindered anticipatory increases in α power reflecting aspecific preparatory stages and dynamic reallocations of attentional resources (van Diepen and Mazaheri, 2017).

A clear modulation of α-band activity was evident shortly after stimulus onset. Alpha desynchronization (Mazaheri and Picton, 2005; Pfurtscheller and Lopes da Silva, 1999) was indeed the main component determining the observed network reconfiguration, accompanied by local increases of α-γ coupling in relevant sensory regions (V5 and MTG for motion discrimination, V1 for orientation discrimination). This latter phenomenon is in line with recent accounts hypothesizing a role of α rhythms in aligning the phase of high-frequency oscillations carrying relevant processing (Bonnefond et al., 2017; Pascucci et al., 2018). Under this view, the decrease of α power resulting from the evoked desynchronization would allow longer windows of neuronal excitability, favoring local increases and feedforward propagation of γ-band activity (Bonnefond et al., 2017). Thus, the co-occurrence of α desynchronization and α-γ coupling may underlie the emergence of short functional windows where the release from inhibitory signals enhances sensory and task-relevant processing. In this respect, our results are also in agreement with the gating-by-inhibition hypothesis that postulates a similar inverse relationship between the degree of α-band synchronization and the level of local neuronal excitability and γ-band activity (Bonnefond and Jensen, 2015; Haegens et al., 2011b; Jensen and Mazaheri, 2010; Mathewson et al., 2011; Mazaheri and Jensen, 2010).

As a means to describe functional network properties, we used graph efficiency measures that indicate how easily signals can travel between network nodes, i.e., the degree of local and global integration, depending on the configuration of directed functional connections. The analysis of network efficiency revealed a sequence of stimulus-evoked brain dynamics involving α-band activity: an initial rapid desynchronization of α rhythms with increased α-γ coupling at the local level (∼50–120 ms post-stimulus), followed by the rise of large-scale γ-band interactions, with enhanced network efficiency in the γ range (150–200 ms post-stimulus). The transition between local and distributed γ-band activity, and the increased efficiency of inter-areal high-frequency communication occurred at latencies that are consistent with stimulus processing time and early attentional effects (Figure 1) (Hillyard et al., 1998; Hillyard and Anllo-Vento, 1998), thus, they may reflect the dynamics by which attentional selection renders stimulus content accessible to widespread circuits.

Our results also suggest distinct mechanisms for general and task-specific attentional effects. General effects were mediated by pre-stimulus changes in spectral power across cortical regions and by topological changes in the brain-wide network of functional interactions. These included increased spectral power in the β-band over lateral-occipital areas and increased network efficiency in the β and γ-band. The results therefore point to an aspecific enhancement of large-scale network communication that facilitates information routing while preparing for an attended stimulus to appear (Buschman and Kastner, 2015; Fries, 2015). After stimulus onset, we found that attention induced desynchronization first in the α and then in β-band (Klimesch et al., 2007, 2001; Mazaheri and Picton, 2005; Pascucci et al., 2018; Schmiedt et al., 2014). These stimulus-evoked effects co-occurred with a sequence of frequency-specific changes in network topology, starting with enhanced network efficiency in the γ-band (∼150–200 ms), followed by reduced efficiency in the α and β-band (after 240 ms). This later decrease of network integration may suggest a return to more local and sparse computations after the critical time of attention-mediated processing.

Task-specific attentional modulations involved more local phenomena, characterized by increased coupling between lower frequencies and γ-band activity. These effects were selective for functionally specialized regions (in V5-MT for motion, in V1 for orientation) during both pre- and post-stimulus time: task-relevant regions showed both anticipatory increases in β-γ coupling and reactive increases in α-γ coupling. Based on the well-known specialization of these regions in processing the two types of stimulus’ features used (Ahlfors et al., 1999; Bosking et al., 1997; Koelewijn et al., 2011; Schoenfeld et al., 2007; Simoncelli and Olshausen, 2001), these results support the view of PAC and nested oscillations as regulatory mechanisms for the local analysis and feedforward communication of relevant sensory signals (Jensen and Mazaheri, 2010; Pascucci et al., 2018).

In this work, we provide first evidence of a sequential relationship between anticipatory increases in β-γ PAC and stimulus-evoked α-γ coupling. These two local phenomena may be linked to precise endogenous and exogenous components of attentional processing: pre-stimulus anticipatory effects could promote the endogenous reactivation of task-specific content in sensory circuits (Spitzer and Haegens, 2017), whereas post-stimulus coupling may reflect the enhanced processing of relevant exogenous signals (e.g., target stimuli) (Bonnefond and Jensen, 2015; Jensen and Mazaheri, 2010). Theories of communication based on nested oscillations propose that low-frequency carriers in the theta (θ, 3–7 Hz), α or β-band establish inter-areal communication at larger scales, by mediating local excitability (reflected by γ-band activity) (Bonnefond et al., 2017). Cross-frequency coupling at the local level, therefore, would coordinate γ-band activity and information routing at different scales (Buzsáki, 2006; Buzsáki and Wang, 2012; Canolty et al., 2006; Canolty and Knight, 2010; Jensen and Colgin, 2007; Penny et al., 2008; Voytek et al., 2010). Here we show that local integration and network efficiency at the carrier frequency can be also a determinant of local coupling and information flow. This was evident in the task-specific anticipatory increase of β-band local efficiency, which showed a positive relationship with the degree of β-γ PAC enhancement due to attention. Stimulus-evoked α-γ PAC, however, did not correlate with network properties, but was instead driven solely by α-band desynchronization. This indicates a different mechanism that may emerge upon the release from inhibition. Adding on the gating-by-inhibition framework (Mazaheri and Jensen, 2010), we found that the effects of the release from inhibitory gating are selective to the upcoming tasks, occurring exclusively in cortical regions encoding task-relevant features. A compelling question for future investigations is whether these distinct mechanisms of local cross-frequency coupling (anticipatory β-driven, reactive α-driven) underlie also the attentional selection based on whole-object representations, for example of faces and houses via selective involvement of the fusiform face area and parahippocampal place area, respectively.

For functional connectivity analyses, we employed state-of-the-art multivariate methods (Baccalá and Sameshima, 2014), and a recently developed adaptive filter that was specifically designed for fast-changing signals like VEPs (Pascucci et al., 2019). While we considered a large-scale network of 22 brain-wide cortical regions for our analyses, one potential limitation is linked to unobserved common inputs (Bastos and Schoffelen, 2016; Pagnotta et al., 2018a). Previous studies revealed that the pulvinar and mediodorsal thalamus play a role in attentional control in non-human primates (Fiebelkorn et al., 2019; Saalmann et al., 2012). We did not include the thalamus as a ROI, due to the EEG intrinsic limitations in accurately reconstructing subcortical activity (although see Seeber et al., 2019). Hence, our results provide solely an account for the cortico-cortical interactions in mediating selective attention. In the future, the use of invasive recordings in clinical populations and implanted patients may provide further insights about the role of subcortical structures and thalamo-cortical interactions in controlling attention and other cognitive functions.

## 4. Methods

### 4.1. Participants

Twenty healthy human participants (13 female; all right-handed; ages 19–34 y, *M*=23.25, *SD*=4.15) with normal or corrected-to-normal vision (visual acuity across participants 1.10–1.63, *M*=1.40, *SD*=0.19; Freiburg visual acuity test (Bach, 1996)) took part in the study for monetary compensation (20.– CHF/hour). The study was performed in accordance with the Declaration of Helsinki on “Medical Research Involving Human Subjects” and after approval by the responsible ethics committee (Commission cantonale d’éthique de la recherche sur l’être humain, CER-VD). Written informed consent was obtained from each participant prior to the experimental sessions. Data from one participant (male, age = 22) were excluded because of excessive artifacts in the EEG.

### 4.2. Experimental design

Visual stimuli were Random Dot Kinematograms (RDK: 10° field-size, 1200 dots, 0.2° dot-size, infinite dot-life, and 4°/s dot-speed), that were contrast-modulated through a Gaussian-windowed sinusoidal grating (Gabor: 6° width at 3 SDs, and 0.5 cycles/° spatial frequency), such that the Gabor determined the visible region of the RDK and its spatial pattern of contrast (Figure 1A). The orientation of the Gabor ranged between 45° and −45° off-vertical, and phase varied randomly at every trial. In the RDK, a subset of dots moved coherently either towards left or towards right (signal-dots), while the remaining dots moved with random walks (noise-dots), in such a way that their direction varied at every frame but their speed was kept constant. Stimuli were presented on a gray background for 300 ms always at the same spatial location, around a central fixation spot that was constantly on the screen (0.2° size, and (0,0,160) color in RGB digital 8-bit notation). The inter-trial interval (ITI) was randomly varied between 800 and 1000 ms. The generation and presentation of the visual stimuli was performed using a combination of in-house Python codes and PsychoPy Builder (Peirce et al., 2019).

The study consisted of three separate experimental sessions: one behavioral (approx. 45 minutes), one EEG (approx. 60 minutes), and one session of fMRI (approx. 45 minutes). In all sessions, the participants were asked to performed different tasks on the same type of visual stimuli. The behavioral session comprised two different tasks in separate blocks (100 trials), requiring a two-choice response. At the beginning of each block, participants were either instructed to report the perceived motion direction in the RDK (left vs. right, motion discrimination task), or the off-vertical tilt of the Gabor (left vs. right, orientation discrimination task), by pressing the corresponding key on a keyboard. The aim of the behavioral session was to determine individual thresholds for coherent motion and orientation discrimination, in order to level performance for the two tasks in the EEG and fMRI sessions, keeping accuracy at 82% of correct responses for both tasks. Thresholds were estimated using an adaptive staircase procedure (Watson and Pelli, 1983). We used starting values of 60% for signal-dots and 4° for angle of orientation, while the task-irrelevant feature was kept constant during the staircase procedure (40% of signal-dots in the orientation discrimination task and ±3° off-vertical in the motion discrimination task). This allowed to obtain individual discrimination thresholds at the same level of accuracy across participants, both for the motion discrimination task (signal-dots percentages 10.61–89.82, *M*=44.50, *SD*=29.09) and for the orientation discrimination task (angles of orientation 1.53°–4.30°, *M*=2.45°, *SD*=0.72°). These individually calibrated values were used to define stimuli in the subsequent EEG and fMRI sessions.

During neuroimaging sessions, participants also performed an additional control task, where they had to report sporadic color changes in the fixation spot lasting 200 ms (from dark blue to red, 30% of trials; Figure 1A) by pressing a response button. To ensure that the timing of these color changes was unpredictable, we presented them at any time during the trial, randomly. Trials in which the color change overlapped with visual stimuli appearance were excluded from the successive analyses. This design resulted in two conditions in which the RDK was task-relevant (motion and orientation discrimination) and one condition where it was task-irrelevant (color change detection). We refer to these different task conditions as Attended (Attended-motion and Attended-orientation) and Unattended, indicating the type of attentional processing required for the visual stimuli. The EEG session consisted of 160 trials divided in 4 blocks for each of task condition (12 blocks in total). Small breaks were offered between blocks. The fMRI session followed a similar block structure, but with less trials (48 trials per task condition). A fixed 12 s break was presented after each block, following the instruction to “REST” (Rest period), with the exception of the last Rest period that lasted 60 s. Task order was counterbalanced across participants in each of the three experimental sessions. Participants had a limited time to respond (1500 ms), after which the response for that trial was considered incorrect.

### 4.3. MRI materials and procedures

#### 4.3.1. Data acquisition and preprocessing

MRI data were collected using a Discovery MR750 3.0T (GE Healthcare, Chicago, USA) at the cantonal hospital, HFR Fribourg. First, functional data were collected with T2*-weighted EPI sequence with 40 slices each, 3 mm slice-thickness, 0.3 mm spacing between slices, interleaved bottom-up slice acquisition, anterior-to-posterior phase encoding direction, 2500 ms repetition rime (TR), 30 ms echo time (TE), and 85° flip-angle. The first 4 volumes of each run were discarded from analyses. Functional data acquisition was performed while each participant performed the visual discrimination tasks. Behavioral responses were collected using an MRI-compatible fiber optic response pad (Current Designs Inc., Philadelphia, USA). Stimuli were presented on a NordicNeuroLab (Bergen, NO) MRI-compatible LCD monitor (32 inches diagonal size, 1920 × 1080 resolution, 405 c/m^2^ surface luminance, 4000:1 contrast, 60 Hz refresh rate, 6.5 ms response time), placed above the scanner-bed at a distance of 244 cm from the participant’s eyes and visible to the participant through a mirror placed on the head coil. Participants who needed optical correction (e.g., for myopia) wore MRI-compatible glasses with the appropriate lenses. Next, an anatomical whole-head image was acquired using a T1-weighted BRAVO 3D sequence with 280 slices, 1 mm isotropic voxels, 730 ms TR, 2.8 ms TE, 9° flip-angle, 256 x 256 mm^2^ field of view (FOV), and 900 ms inversion time (TI).

The preprocessing of both functional and structural images was performed using the Statistical Parametric Mapping (SPM) toolbox (Penny et al., 2011), in its SPM12 version (University College London, London, UK; https://www.fil.ion.ucl.ac.uk/spm/). Functional images were first aligned to the mean of each session using a two-pass realignment procedure for motion correction, and successively a slice-timing correction was applied (Sladky et al., 2011). After realignment, the mean functional image was coregistered to the anatomical image using the normalized mutual information as cost function. The standard segmentation procedure in SPM12 was applied to obtain individual masks for cerebrospinal fluid (CSF) and white matter (WM), which were used to extract the time courses of CSF and WM signals for each participant. At last, all images were normalized to the Montreal Neurological Institute and Hospital (MNI) stereotaxic space using a fourth-order B-spline interpolation, and smoothed with a Gaussian filter (8 mm FWHM kernel).

#### 4.3.2. fMRI statistical analysis

A two-stage approach based on a general linear model (GLM) was employed to analyze the functional images (Friston et al., 1994). In this approach, the first-level analysis was implemented using a block design with four regressors of interest, each modeled with a boxcar function convolved with the canonical hemodynamic response function (HRF). Four regressors of interest were defined to model the two Attended conditions (Attended-motion and Attended-orientation), the Unattended condition, and Rest periods. The GLM included also a set of nuisance regressors that modeled the six motion realignment parameters, the mean signals in CSF and WM, and a constant term. Finally, a high-pass filter (200 s cutoff) was applied to the functional images time series, which allowed removing noise at very low frequencies. At the first-level individual analysis, two contrasts were considered to obtain task-related activation maps: i) “Attended vs. Unattended”; ii) “Attended vs. Rest”. The difference between the two Attended conditions and the Unattended was computed for the first contrast, which allowed identifying areas that were differentially active across task-relevant and task-irrelevant conditions (two-tailed test). The second contrast was based on the difference between Attended conditions and Rest period, which allowed identifying areas more strongly engaged by the tasks (one-tailed test). The second-level group analysis was implemented on the previously obtained statistical maps and involved voxel-wise *t*-test comparison across participants. Here, participants were modeled as random effects, and statistical significance was assessed at the group-level using an uncorrected voxel-based threshold *p*<0.001, together with a minimum cluster-size of 5 voxels (Figure 3A–B). The results of this procedure were used to extract ROIs for source-space EEG analyses (see below).

### 4.4. EEG materials and procedures

#### 4.4.1. Data acquisition and preprocessing

EEG recordings were collected at the Department of Psychology, University of Fribourg. During the EEG session, participants were sitting inside a dark shielded room, with their head leaning on a chinrest positioned at 71 cm from a VIEWPixx /EEG™ (VPixx Technologies Inc., Saint-Bruno, CA) LCD monitor (24 inches diagonal size, 1920 x 1080 resolution, 100 cd/m^2^ luminance, 120 Hz refresh rate, 1 ms pixel response time). A 2-button RESPONSEPixx response box (VPixx Technologies Inc.) was plugged into the monitor and used by the participants during the visual tasks. EEG data were acquired using a 128-channel ActiveTwo EEG system (Biosemi, Amsterdam, NL). Data acquisition was performed at a sampling rate of 1024 Hz, and signals were referenced to the Common Mode Sense (CMS) active electrode, which together with the Driven Right Leg (DRL) passive electrode formed a feedback loop in the system. This CMS/DRL loop allow to drive the common-mode voltage as close as possible to the ADC reference voltage in the AD-box, and to obtain an extra 40 dB common-mode rejection ratio at 50 Hz, when compared with using normal ground electrodes with same impedance. At the end of the EEG session, the 3D coordinates of the positions of the electrodes were localized for each participant using an ELPOS system (Zebris Medical GmbH, Isny im Allgäu, DE). These individual electrodes’ positions were used for EEG source reconstruction procedure (see section 4.5).

EEG preprocessing was performed using a combination of EEGLAB (Delorme et al., 2011; Delorme and Makeig, 2004), its plugins, and in-house scripts implemented in MATLAB (The MathWorks Inc., Natick, USA). Data were first downsampled to 500 Hz using an anti-aliasing filter with 125 Hz cutoff frequency and 50 Hz transition bandwidth. Afterwards, the data were detrended using high-pass filtering (1 Hz low-frequency cutoff), as implemented in the PREP plugin (Bigdely-Shamlo et al., 2015). Line noise and its harmonics were reduced using the adaptive filtering technique implemented in the EEGLAB plugin CleanLine (https://www.nitrc.org/projects/cleanline), which allows identifying and removing significant sinusoidal artifacts. Epochs were extracted using the time window −1500– 1000 ms around stimulus onset. Noisy EEG channels were identified by visual inspection and removed before proceeding to the subsequent preprocessing steps (number of removed channels across participants 12–21, *M*=15.85, *SD*=2.64). Epochs contaminated by noise artifacts and eye movements artifacts, e.g., eye blinks occurring within 500 ms from stimulus presentation (either before or after), were also rejected by visual inspection (percentage of rejected epochs across participants 12.50– 38.95%, *M*=23.01, *SD*=7.49). Decomposition of EEG data by independent component analysis (ICA) was performed using the FastICA algorithm (Hyvärinen and Oja, 2000). ICA components identified as eye artifacts or muscular activity artifacts were removed from the data (number of removed components across participants 5–23, *M*=10.30, *SD*=4.09). As final preprocessing steps, noisy EEG channels were interpolated using a spherical spline interpolation (Perrin et al., 1989); then, the signals were re-referenced to the common average reference (Lehmann and Skrandies, 1980). We excluded trials with incorrect responses, and the amount of trials across task conditions was balanced within-subjects (number of trials per task condition across participants 74–127, *M*=103.5, *SD*=15.3). This choice was motivated by the fact that functional connectivity analysis can be sensitive to the amount of trials (Astolfi et al., 2008; Toppi et al., 2012), and consequently an imbalance of trials between task conditions may create spurious differences from comparing them.

#### 4.4.2. Visual-evoked potentials and power spectra

The VEPs of each participant were obtained by separately averaging the epochs of the three task conditions. VEPs analysis was confined to three clusters on sensor-space: centro-occipital (CO), left occipito-temporal (lOT) and right occipito-temporal (rOT). Each electrode-cluster comprised 7 electrode sites from the 128-channel Biosemi cap, using the same subdivision adopted in Daffner et al. (2012). Each trial was baseline corrected before computing VEPs, using the 200 ms pre-stimulus interval as baseline. A repeated-measures ANOVA was employed to compare the three task conditions in each of the pre-selected sensor-space macro-area. Statistical significance was assessed at alpha-level of 0.05, using false discovery rate (FDR) correction for multiple comparisons (Benjamini and Hochberg, 1995). In addition, a temporal stability criterion was imposed on the results so that only significant results that exhibited stability over at least 5 time frames (10 ms) survived. Post-hoc analysis was carried out using the Fisher–Hayter procedure (Hayter, 1986), in each of the time intervals that showed a statistically significant difference from the repeated-measures ANOVA.

Power spectral density analysis was performed on sensor-space time series in two time windows of interest: pre-stimulus (anticipatory) and post-stimulus (reactive). The anticipatory window was defined as the last 300 ms before stimulus onset. The reactive window was selected as the interval 200–500 ms after stimulus onset, because it is when strong stimulus-induced power changes are typically observed at low-frequencies (Klimesch et al., 2007, 2001; Mazaheri and Picton, 2005; Schmiedt et al., 2014). Power spectrum at each electrode location was estimated in the frequency range 1–100 Hz, using a multitaper method based on orthogonal tapers given by Slepian sequences, whose complete description and MATLAB implementation were provided in previous works (Mitra and Pesaran, 1999; Pagnotta et al., 2018a, 2018b). In this approach, the time-bandwidth product (*NW*) regulates the trade-off between variance and bias of the spectral estimates. To accommodate for the different characteristics of the spectral components at low and high frequencies, two distinct time-bandwidth products were here used for frequencies up to 45 Hz (*NW*=3) and for frequencies above 45 Hz (*NW*=16). Attended and Unattended conditions were compared by using a cluster-based permutation approach over frequencies and EEG electrodes locations, with 30 mm maximum distance to define neighboring electrodes. In this and every other analysis using a cluster-based permutation approach, to compare Attended and Unattended conditions, we employed the same settings: two-tailed *t*-test (*p*<0.05), 50000 permutations, and *p*<0.05 for the permutation test (Maris and Oostenveld, 2007).

### 4.5. EEG source reconstruction

Source reconstruction techniques enable to estimate and localize the EEG sources of brain electrical activity, by solving first a forward problem and then an inverse problem (Michel et al., 2004). For the forward problem, the volume conduction model of each participant was constructed using a boundary element method (BEM) (Hamalainen and Sarvas, 1989), which employed the information about boarder surfaces between three different tissue-types: scalp, skull and brain. These surfaces were obtained from a segmentation procedure of the individual-participant anatomical MRI, using a Gaussian kernel for smoothing (5 voxels FWHM). The participant-specific 3D electrode coordinates (see section 4.4.1) were aligned to the head model of the same participant using an interactive re-alignment procedure, followed by a projection of electrodes onto the head surface. For group analyses on source-reconstructed EEG data, we employed a template grid based on MNI template anatomical MRI to define the solution points. The template comprised 1725 source-points (equivalent current dipoles) with 10 mm spacing, and was constrained to be within the cortical gray matter. Individual MRIs were warped to the template, and the inverse of this warping procedure was applied to the template grid. This procedure provided participant-specific grids that were no longer regularly spaced, but that became equivalent across participants in normalized MNI space, facilitating group analysis on source-space. Finally, individual lead field matrices were computed considering an unconstrained-orientation forward operator, i.e., each solution point was modeled as three orthogonal equivalent current dipoles placed at that location.

To solve the EEG inverse problem, we employed the linearly constrained minimum variance (LCMV) beamformer (Van Veen et al., 1997). The LCMV belongs to the family of spatial filtering methods and its implementation relies on the use on an estimate of the sensor-space covariance matrix, which was here estimated from the time window −500–500 ms around stimulus onset. Source-reconstructed signals were extracted from 22 cortical ROIs using an approach based on singular value decomposition (SVD), estimating scalar-value time series that explain most of the variability across dipoles in each ROI (Rubega et al., 2019). The ROIs were defined with an automatic procedure, starting from the spatially segregated cluster-peaks obtained from the fMRI statistical analysis (Figure 3A–B). First, each fMRI cluster-peak was used as the center of a sphere of 15 mm radius. Then, all solutions points inside the sphere were selected, and tissue-type consistency was verified using the automated anatomical labeling (AAL2) atlas parcellations defined in MNI space, so that only the solution points that fell within the atlas areas were kept. This procedure allowed us to uniquely identify each ROI, in such a way that no solution points were shared across ROIs. Table 2 lists the 22 ROIs (Figure 3C–E). As final step, an approach based on innovations orthogonalization was used for leakage correction (Pascual-Marqui et al., 2017). This orthogonalization approach allowed reducing the detrimental effects on functional connectivity analyses of zero-lag cross-correlations, which are due to instantaneous linear mixing between source-reconstructed signals (source leakage) and are known to produce spurious functional connectivity estimates (Anzolin et al., 2019). All the steps of EEG source reconstruction were implemented using in-house MATLAB codes and routines from FieldTrip (Oostenveld et al., 2011) (Radboud University, Nijmegen, NL; http://www.ru.nl/neuroimaging/fieldtrip).

### 4.6. Time-varying power spectra

To investigate local brain rhythms modulations depending on whether the stimuli were relevant or not, time-varying power spectra were estimated from source-reconstructed signals of all ROIs using a Morlet wavelet transform with central frequency parameter *ω*_*0*_ = 6, in combination with zero-padding to solve the problem of edge effects (Torrence and Compo, 1998). Time-varying spectral estimates were estimated in the frequency range 1–100 Hz. This procedure was performed for each participant and task condition using the MATLAB codes made available in Pagnotta et al. (2018b), after subtracting the ensemble mean across trials from single-trial data, which allows minimizing the influence of evoked responses on power spectra (Kalcher and Pfurtscheller, 1995; Rajan et al., 2018). A cluster-based permutation approach was used to compare time-varying power spectra of Attended and Unattended conditions, over time frames (−50–600 ms around stimulus onset) and frequencies (1– 100 Hz).

### 4.7. Local cross-frequency coupling

#### 4.7.1. Anticipatory (pre-stimulus) cross-frequency coupling

Cross-frequency coupling was measured in terms of dependence between phase of low-frequency oscillations and amplitude of high-frequency oscillations, called phase-amplitude coupling or PAC. Among the available types of cross-frequency coupling, we employed the PAC because of its more clear functional role and physiological plausibility (Canolty and Knight, 2010; Voytek et al., 2010). Several approaches have been proposed to measure PAC (Canolty et al., 2006; Martínez-Cancino et al., 2019; Penny et al., 2008; Tort et al., 2010; Voytek et al., 2013). We here employed the modulation index based on GLM (Penny et al., 2008) to calculate within-region PAC before stimulus onset, using the toolbox from Martínez-Cancino et al. (2019). The ensemble mean was preliminary subtracted from single-trial data for each participant and task condition before PAC estimation. The analysis was restricted to the subset of 9 occipito-temporal ROIs. The central frequency for the phase time series (*f*_*PHASE*_) ranged from 4 Hz to 30 Hz in 1Hz-steps, and the higher central frequency for the amplitude time series (*f*_*AMPLITUDE*_) ranged from 40 Hz to 100 Hz in 2Hz-steps. The bandwidth of the band-pass filters to estimate the phase time series (*BW*_*PHASE*_) and the amplitude time series (*BW*_*AMPLITUDE*_) were defined according to the following expressions:

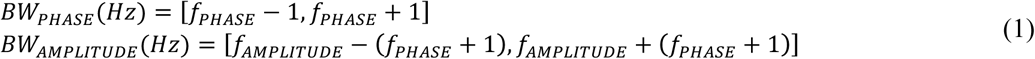

Filtering was applied to the pre-stimulus interval (−300–0 ms), using two buffer windows of 1850 ms, before and after the interval of interest. Each buffer window consisted of 600 ms of observed signal and the rest zeros, such that filtering was applied every time to a 4000 ms window. A combination of high-pass and low-pass finite impulse response (FIR) filters was employed to band-pass filter the data in the frequency bands of interest. Next, the Hilbert transform was applied to extract the analytic signals from which instantaneous low-frequency phase and high-frequency amplitude time series were estimated, which allowed computing PAC in the time window of interest using the GLM-based method. This procedure was repeated for every combination of *f*_*PHASE*_ and *f*_*AMPLITUDE*_, in each trial. The averages across trials of PAC estimates were compared over all pairs of *f*_*PHASE*_ and *f*_*AMPLITUDE*_ between Attended and Unattended conditions, using a cluster-based permutation approach. We excluded from the analysis all the *f*_*PHASE*_–*f*_*AMPLITUDE*_ pairs for which *BW*_*AMPLITUDE*_ overlapped with *BW*_*PHASE*_.

Since strong imbalances in the power spectra between conditions can lead to spurious PAC differences (Aru et al., 2015), a control analysis based on a stratification procedure was performed separately for each ROI using the *ft_stratify* function from FieldTrip (Oostenveld et al., 2011), with 1000 iterations. At each iteration, this procedure selects subsets of trials from the two conditions with matched distributions of power spectra across trials, for both phase and amplitude frequency bands. Balanced numbers of trials between conditions were maintained during this procedure. The final estimates from this control analysis were obtained as the median across iterations of the power-balanced estimates, and these were compared between Attended and Unattended conditions using the cluster-based permutation. The control analysis based on stratification was used to confirm the between-condition differences in pre-stimulus PAC.

#### 4.7.2. Reactive (post-stimulus) cross-frequency coupling

A measure of stimulus-evoked cross-frequency coupling was computed for each task condition using the instantaneous phase-locking value (PLV) between the low-frequency phase and the phase of amplitude-filtered high-frequency signal (Lachaux et al., 1999; Pascucci et al., 2018). As in the pre-stimulus analysis, the ensemble mean was preliminary subtracted from single-trial data for each participant and task condition, and the estimation was performed on the 9 occipito-temporal ROIs. Time-varying PLV estimates were obtained at each time frame by averaging the instantaneous contributes across trials. The analysis was performed on the interval 0–500 ms after stimulus onset, using two buffer windows (before/after), each of 1450 ms length (including 300 ms of observed signal). Low-frequency ranges were separately defined in α and β-band, as *f*_*PHASE*_ = 10 Hz and *f*_*PHASE*_ = 20 Hz for the comparison between Attended-motion and Unattended, and as *f*_*PHASE*_ = 8 Hz and *f*_*PHASE*_ = 17 Hz for the comparison between Attended-orientation and Unattended. These values were selected in data-driven way, on the basis of the task-specific results obtained by comparing time-varying power spectra (Figure 2D and Figure S1D). For the amplitude time series, *f*_*AMPLITUDE*_ ranged from 50 Hz to 100 Hz in 2Hz-steps. The bandwidths of the two band-pass filters were chosen using the expressions in equation (1) (see section 4.7.1). The procedure of filtering (FIR-based) and analytic signals extraction (using Hilbert transform) was carried out as previously done for the anticipatory PAC analysis. A cluster-based permutation approach over time frames and amplitude high-frequencies was employed to compare PLV estimates between Attended and Unattended conditions.

### 4.8. Large-scale network analyses

To estimate the functional interactions among brain regions we used the *information* form of partial directed coherence (iPDC) (Takahashi et al., 2010). The iPDC is based on the notion of Granger-Geweke causality (Geweke, 1984; Granger, 1969), which relies on the concepts of temporal precedence and statistical predictability between simultaneously recorded time-series, extended to the spectral-domain (Seth et al., 2015). Computationally the iPDC is derived from a multivariate autoregressive (MVAR) model with a certain order, which determines the number of past observations included in the model. To derive iPDC estimates the MVAR coefficients are Fourier transformed, appropriately weighted by noise covariance matrix contributes to overcome the scale-variance problem, and finally normalized (Baccalá and Sameshima, 2014). The iPDC is a measure of the partialized delayed functional interactions between brain regions, which is able to characterize the interaction directionality between two regions (directed measure), as well as discerning direct from indirect or mediated paths of connections (directness). More precisely, the iPDC from the *j*-th time series to the *i*-th time series is equivalent to the squared coherence between the innovation process associated with *i* and the partialized version of *j*, which via appropriate logarithmic expression and integration provides the mutual information rates between these two processes (Baccalá and Sameshima, 2014; Takahashi et al., 2010). In the present study, the iPDC was used as measure of the frequency-specific, directed connections between ROIs, and all connectivity analyses were performed on source-reconstructed time series, because analyses applied on EEG sensor-space time series do not allow any meaningful interpretation in terms of interacting brain sources (Brunner et al., 2016; Van de Steen et al., 2016).

#### 4.8.1. Anticipatory (pre-stimulus) functional connectivity

To investigate the anticipatory network-level modulations of attention, we performed a directed functional connectivity analysis on the pre-stimulus interval (−300–0 ms). Preprocessing included a signal decimation by a factor of 2 (new sampling rate of 250 Hz), the removal of temporal mean and ensemble mean from single-trial data to meet zero-mean assumptions, and the use of the augmented Dickey–Fuller test for stationarity as implemented in the Oxford MFE Toolbox (https://www.kevinsheppard.com/code/matlab/mfe-toolbox/). The optimal model order for each participant was selected by first identifying the value that minimized the difference between power spectra obtained from parametric MVAR-modeling and those obtained using a nonparametric approach based on multitaper (see section 4.4.2), in each task condition separately, and then selecting the maximum value across task conditions (model orders across participants 7–18, M=14.05, SD=3.27). These individual-participants’ fixed values were then used for analysis. A stepwise least squares MVAR model estimation was then performed on source-reconstructed signals to derive iPDC estimates in the network of 22 ROIs, for each task condition. The iPDC estimates were computed in the frequency range 1–100 Hz (in 1Hz-steps).

Measures from graph theory were used to characterize large-scale topological properties of the frequency-specific, directed functional networks in the different task conditions (Rubinov and Sporns, 2010). Each ROI was considered as a node in the graph and two measures were estimated: global efficiency and local efficiency. The network’s global efficiency is the average inverse shortest path length and indicates the level of global functional integration in the network (Latora and Marchiori, 2001). The single node’s local efficiency is the global efficiency of the local subgraph comprising all its immediate neighbors (Latora and Marchiori, 2001), and the network’s local efficiency is computed as average efficiency of all local subgraphs, which represents the level of local integration within subgraphs. High local efficiency suggests functional segregation within the network (Rubinov and Sporns, 2010). Both global efficiency and local efficiency were computed for each participant and task condition from the full matrix of iPDC estimates, i.e., from a fully-connected, weighted and directed adjacency matrix. Subgraphs and nodes’ neighbors were thus determined by the interaction strengths: the stronger the connection weight between two nodes, the closer they were (functionally).

When we consider global and local efficiency derived from iPDC-based graphs, we can talk about frequency-specific levels of functional integration among brain regions, respectively globally and within subgraphs, for two reasons. First, the iPDC has a clear interpretation in terms of between-processes mutual information rates, providing an association with the notion of information flow (Baccalá and Sameshima, 2014; Takahashi et al., 2010). Second, thanks to its partialization approach, the iPDC provides the directness of functional connections. It is therefore reasonable to characterize the cascades of these direct connections as paths of integration between regions, unlike when using classical measures such as cross-correlations and spectral coherence (Rubinov and Sporns, 2010).

Each graph measure was separately compared between Attended and Unattended conditions (two-tailed *t*-test, *p*<0.05, with FDR correction). Graph measures were derived using the Brain Connectivity Toolbox (Rubinov and Sporns, 2010) (http://www.brain-connectivity-toolbox.net). All the other steps of the functional connectivity analysis were performed using in-house scripts implemented in MATLAB.

#### 4.8.2. Reactive (post-stimulus) functional connectivity

To accommodate the non-stationarity of VEP signals and obtain time-varying functional interactions among brain regions, the iPDC can be derived from a time-varying MVAR (tvMVAR) model of the signals, which can be computed using adaptive algorithms (Arnold et al., 1998; Milde et al., 2010; Pagnotta and Plomp, 2018). This procedure allows obtaining iPDC estimates over frequencies and time frames. Time-varying iPDC estimates were computed in the interval −1500– 800 ms around stimulus onset, for each participant and task condition, after decimating single-trial data by a factor of two and subtracting their temporal and ensemble means. We performed tvMVAR modeling with the Self-Tuning Optimized Kalman filter (STOK) algorithm (Pascucci et al., 2019), using a percentage of variance explained of 99% for setting the filtering factor threshold (Hansen, 1987). Similarly to the anticipatory connectivity analysis, the model order was selected by comparing parametric and nonparametric power spectra in the post-stimulus interval 0–500 ms (model orders across participants 11–21, *M*=17.53, *SD*=2.76).

Measures of time-varying global efficiency and local efficiency were estimated in the time window - 20–500 ms in three frequency bands of interest (α, β, and γ-band). For each frequency band, the time-frequency iPDC estimates were first averaged over frequency bins inside that band, then used to derive time-varying global efficiency and local efficiency, and finally compared between Attended and Unattended conditions (paired sample *t*-test, *p*<0.05 with FDR correction). For the comparison between Attended-motion and Unattended, the frequency bands were selected as 8–12 Hz (α-band), 17–23 Hz (β-band), and 70–95 Hz (γ-band), based on the results on time-varying power spectra (Figure 2D) and α-γ PAC (Figure 6A). For the comparison between Attended-orientation and Unattended, the frequency bands were 6–10 Hz (α-band), 14–20 Hz (β-band), and 65–95 Hz (γ-band) (Figure S1D and Figure 6B).

### 4.9. Correlation between local network properties and PAC

In cortical regions that showed significant anticipatory changes in PAC, we investigated whether there was a relationship between the local PAC and the local network properties of that region, as revealed by the single-node’s local efficiency. In each region, we identified the cluster of *f*_*PHASE*_–*f*_*AMPLITUDE*_ pairs that showed significant differences between Attended and Unattended conditions (Figure 5), which allowed defining also the significant frequency ranges for *f*_*PHASE*_ and *f*_*AMPLITUDE*_. For example, from the comparison between Attended-motion and Unattended in region V5-R the significant cluster is highlighted in opaque red in Figure 5A (β-γ PAC), and the two significant ranges are 24–27 Hz for *f*_*PHASE*_ (β-band) and 54–80 Hz for *f*_*AMPLITUDE*_ (γ-band). Participant-specific PAC estimates were averaged over the significant cluster, separately for Attended-motion and Unattended, and a variation in PAC (*ΔPAC*) was computed as the difference between Attended-motion and Unattended, normalized by the latter. In a similar way, but separately for the two significant ranges for *f*_*PHASE*_ and *f*_*AMPLITUDE*_ (in the example, β-band and γ-band), participant-specific estimates of the region’s local efficiency were averaged over each significant range, separately for Attended-motion and Unattended, and a variation in single-node local efficiency (*ΔE*_*local*_) was computed as the difference between Attended-motion and Unattended, normalized by the latter. We estimated the correlation between *ΔPAC* and *ΔE*_*local*_ across participants using the Spearman’s rank correlation coefficient. Outliers were removed using a robust regression with bisquare weighting function, as implemented in *robustfit* function in MATLAB, on the basis of the studentized residuals (*t*-distribution with 17 degrees of freedom, *p*<0.05). The same analysis was performed for the comparison between Attended-orientation and Unattended, focusing on the set of regions that showed significant anticipatory differences in PAC (Figure 5B).

## Acknowledgements

This study was supported by the Swiss National Science Foundation (grant PP00P1_157420; G.P.). The authors declare that the study has been conducted in the absence of any conflict of interest.

## Supplementary figures

**Figure S1.**
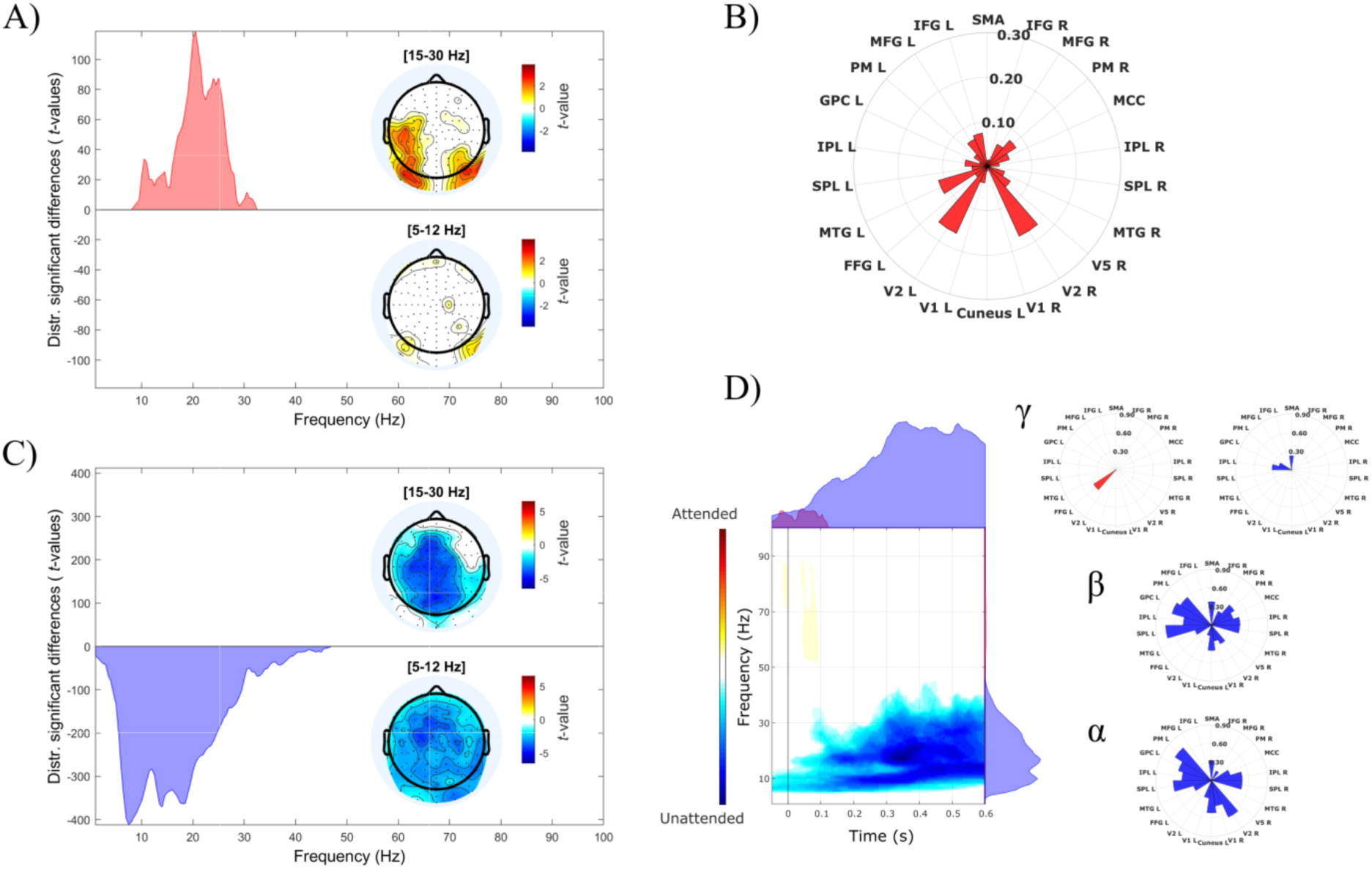
Attentional modulation of brain rhythms: comparison between Attended-orientation and Unattended. Results of sensor-space power spectra analysis are shown for anticipatory (A) and reactive (C) time intervals. Each figure represents the distribution of statistically significant differences (positive *t*-values for Attended-orientation higher than Unattended, and vice versa negative *t*-values) from cluster-based permutation (two-tailed *t*-test with *p*<0.05, 50000 permutations, and *p*<0.05 for the permutation test), together with the topographical distributions of these differences in the α-band (bottom) and β-band (top). B) The polar histogram represents the effect sizes for each ROI of source-space power differences between conditions in the β-band (15–30 Hz). D) The figure shows the results of source-space time-varying power spectra analysis, with the distribution of statistically significant differences from cluster-based permutation (two-tailed *t*-test with *p*<0.05, 50000 permutations, and *p*<0.05 for the permutation test). The marginal plots show time- or frequency-collapsed distributions of significant differences in power spectra. The polar histograms on the right represent the effect sizes for each ROI of power differences between conditions, in the following frequency bands: α (5–12 Hz), β (15–30 Hz), and γ (45–100 Hz). Effect sizes were estimated using Cohen’s *d* (Cohen, 1992).

**Figure S2.**
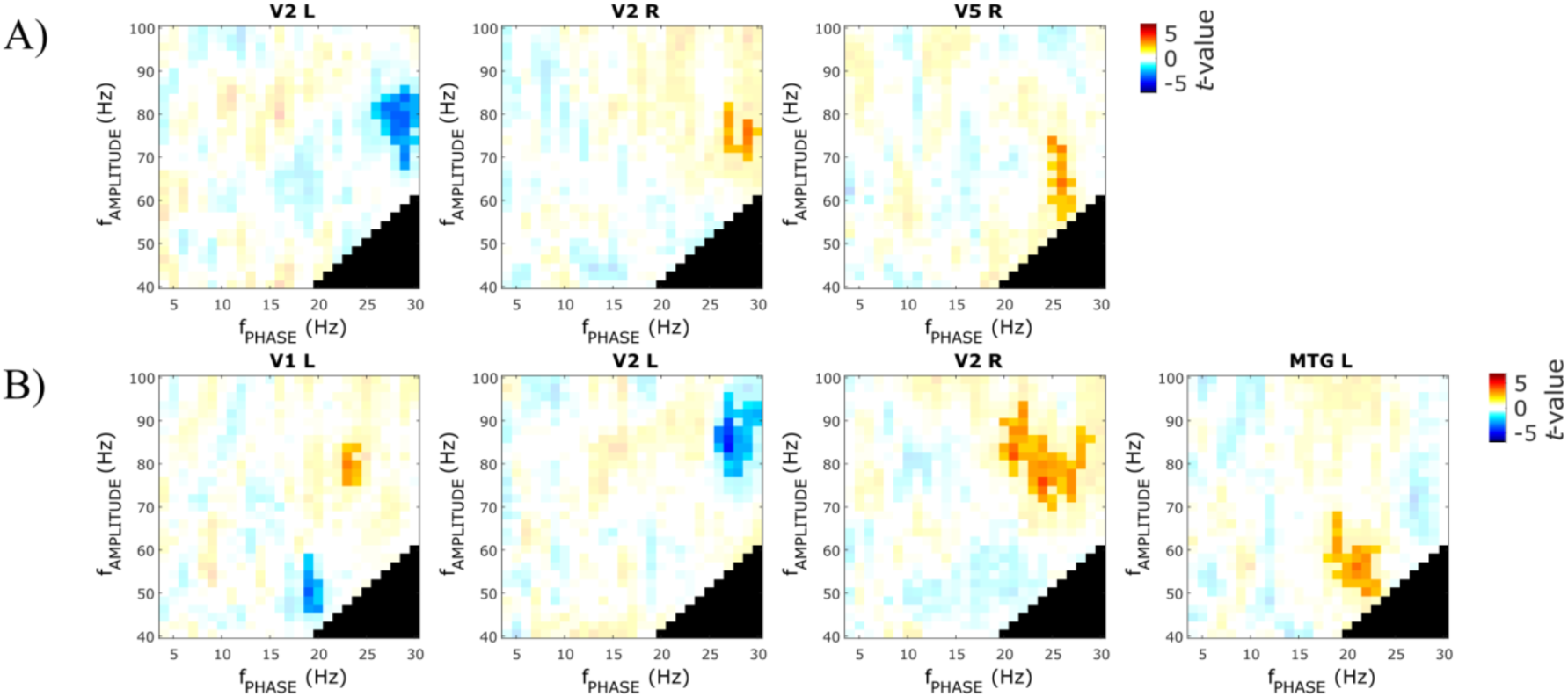
Anticipatory differences in local PAC: control analysis based on stratification (1000 iterations). Differences in within-region anticipatory PAC are shown for each pair *f*_*PHASE*_– *f*_*AMPLITUDE*_, for Attended-motion compared to Unattended (A) and for Attended-orientation compared to Unattended (B). The figure reports only the regions that showed significant differences from the cluster-based permutation comparison (two-tailed *t*-test with *p*<0.05, 50000 permutations, and *p*<0.05 for the permutation test), which are highlighted as more opaque. The black areas indicate *f*_*PHASE*_–*f*_*AMPLITUDE*_ pairs excluded from the analysis (see Methods).

**Figure S3.**
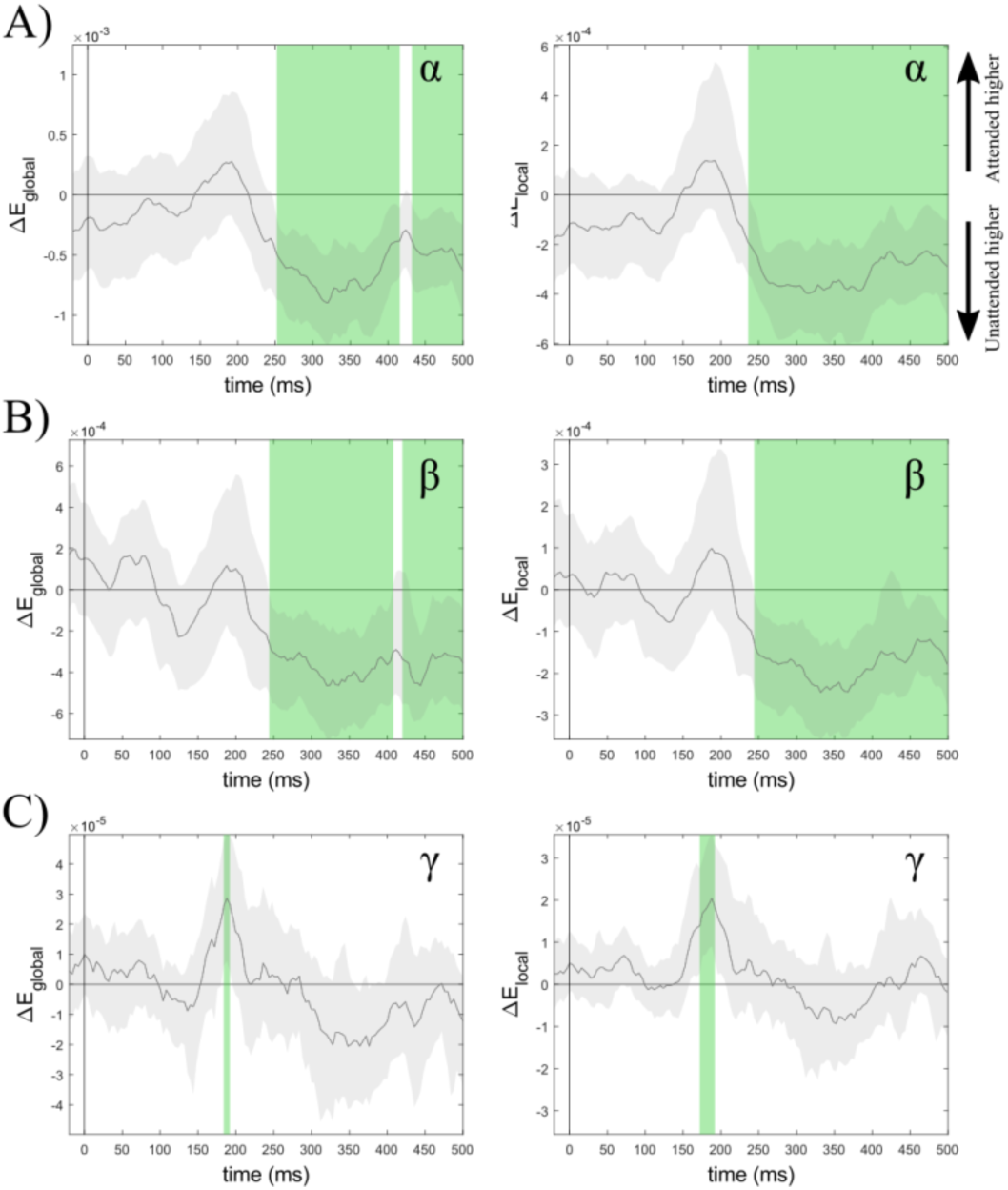
Stimulus-evoked differences in network efficiency. Differences between Attended-orientation and Unattended are shown for three frequency bands, A) α-band, B) β-band, and C) γ-band. Left panels show global network efficiency, right panels local efficiency. Gray shading represents the standard error of the mean. Green vertical shades represent latencies that showed statistically significant results in the two-tailed *t*-test (*p*_*FDR*_<0.05).

